# Determination of *Aedes* mosquito mating success by a rapidly-evolving female-controlled lock-and-key mechanism

**DOI:** 10.1101/2025.04.11.648401

**Authors:** Leah Houri-Zeevi, Madison M. Walker, Jacopo Razzauti, Anurag Sharma, H. Amalia Pasolli, Leslie B. Vosshall

**Author notes:** Correspondence (L.H-Z., Lead Contact), (L.B.V.).

## Abstract

Mosquitoes, the world’s deadliest animal, exemplify single-mating systems where females mate only once in their lifetime, making mate choice critically important for reproductive success and mosquito control. Despite this importance, the mechanisms of female mating control and what prevents additional matings remain poorly understood. To address this gap, we developed a dual-color fluorescent sperm system in invasive *Aedes aegypti* mosquitoes and quantified mating patterns, confirming that 86-96% of females mate only once. Using behavioral tracking of mating pairs, deep learning, and quantitative analysis at increasing resolution, we discovered that females actively control mating initiation through a previously undescribed behavior: genital tip elongation. This female response is triggered by rapidly evolving male genital structures, creating a precise lock-and-key mechanism that determines mating success. Comparative analysis revealed that *Aedes albopictus*, separated from *Aedes aegypti* by 35 million years of evolution, employs a similar female-controlled system. However, *Aedes albopictus* males uniquely bypass female control when attempting cross-species matings with *Aedes aegypti* females but not with conspecific females. This “lock-picking” ability, combined with the known sterility of cross-species matings, may explain how *Aedes albopictus* competitively displaces *Aedes aegypti* populations in overlapping territories. Our findings redefine mosquito reproduction as a female-controlled process and establish a quantitative framework for investigating the molecular and neurobiological mechanisms underlying mating control and species competition in these globally important disease vectors.

## Introduction

Female evaluation and eventual choice of sexual partners shape evolutionary trajectories at multiple levels. These powerful selective forces can drive rapid changes to genomes and genetic programs, nervous systems, morphologies, behaviors, and more^1^. The act of mating is the ultimate product of these processes, with various factors and mechanisms operating together to ensure correct mate choice and successful transfer of genetic material. Females who mate with an unfit male or a male of the wrong species endanger their lineage.

In the highly invasive yellow fever mosquito (*Aedes aegypti*) and Asian tiger mosquito (*Aedes albopictus*), female choice is a high-stakes decision because the female typically mates only once during her lifetime^2–5^. Following this single mating, the female actively stores and nourishes the sperm in internal sperm reservoirs known as *spermathecae*. Females draw on the spermathecae for sperm to fertilize eggs over multiple cycles of blood-feeding and egg production for the rest of their lives^6–8^. With a single and crucial mating event in a lifetime, one might expect elaborate and intricate courtship and mating rituals that would ensure the female correctly assesses and actively chooses her mate.

Such female choice would allow her to block mating attempts from unsuitable males, particularly those of another species^9^. Instead, mating in *Aedes aegypti* mosquitoes typically lasts for 14 seconds and seemingly involves minimal courtship^10^, with the female described to be mostly passive and exerting little if any control over the process^11^. In previous studies, females were observed to kick males as a form of resistance, but this behavior was shown by both virgin and previously mated females^10^. It has been suggested that male and female mosquitoes harmonize their wingbeat as a form of courtship^12,13^, but more recent work has challenged the existence of such a mechanism^14^. Together, these observations create a conceptual tension: the female’s mating decision is crucial, but she is thought to have no control over the process.

*Aedes aegypti* and *Aedes albopictus* are globally invasive mosquito species that transmit dangerous arboviruses to hundreds of millions of people every year^15,16^. While *Aedes aegypti* was first introduced into the New World ∼500 years ago likely through the Atlantic Slave Trade^17^, *Aedes albopictus* only arrived in the last 50 years, and the two species now share broad territories in the Americas^18^. These two *Aedes* species are separated by ∼35 million years of evolution^19^, were geographically separated throughout most of the evolutionary history, and cross-species mating produces no viable progeny^20^. Despite their genetic incompatibility, cross-species mating is observed in wild populations, with an unexplained bias toward *Aedes albopictus* males mating with *Aedes aegypti* females rather than the reciprocal cross-species mating^21^. *Aedes albopictus* males that mate with *Aedes aegypti* females induce a remating block in those females that prevents them from mating with males of their own species, rendering them sterile^21,22^. This mode of reproductive interference by cross-species mating is termed *satyrization*^23–25^. Satyrization is hypothesized to contribute to the ability of *Aedes albopictus* to outcompete *Aedes aegypti* mosquitoes in the southeastern United States, where they share the same territory, leading to observed local extinction of *Aedes aegypti* ^24,26^. The basis for the observed species asymmetry in cross-species mating frequency and satyrization is not known.

In this study, we investigated the behavioral sequence leading to mating in mosquitoes and the mechanisms controlling cross-species mating. We report the first observation of a genital behavior of the female, *female genital tip elongation*, that controls mating in mosquitoes. Only virgin females elongate their genital tips in response to male genital contact. In *Aedes aegypti*, the female genital response is triggered by rapidly evolving secondary male genital structures called *gonostyli*. Fine surgical manipulations of gonostyli dramatically reduced insemination success. Female genital tip elongation also controls mating in *Aedes albopictus* and we identified key differences in genital behaviors and interactions in the two species. Strikingly, we found that *Aedes albopictus* males can bypass female genital tip elongation of both virgin and previously mated *Aedes aegypti* females and initiate mating without the female genital response. These findings correct the long-standing view that the female mosquito plays a passive role in mating and suggest specific mechanisms for reproductive interference by an invasive species that “picks the lock” of the female of the other species.

## Results

### The female remating block is established rapidly

*Aedes aegypti* males pester females and attempt to copulate with them regardless of whether females are virgins or previously mated (**Figure 1A,B**). Because mosquito copulation is rapid^27^, it is not possible to know if these high intensity pestering events and copulation attempts result in insemination, and whether a previously mated female will remate under laboratory conditions. To accurately assess the extent of remating, we generated two transgenic mosquito strains in which sperm express either green fluorescent protein (mNeonGreen) or red fluorescent protein (DsRed2) (**Figure 1C**) under control of the β2Tubulin promoter, which confers expression of the fluorescent proteins in mature sperm^28^. Males from these strains were indistinguishable from wild-type males in their ability to inseminate females (**Figure S1A,B**), and females mated to either transgenic strain produced the same number of eggs and resulted in equivalent larval hatch rates compared to females mated with wild-type males (**Figure S1C-E**).

**Figure 1:**
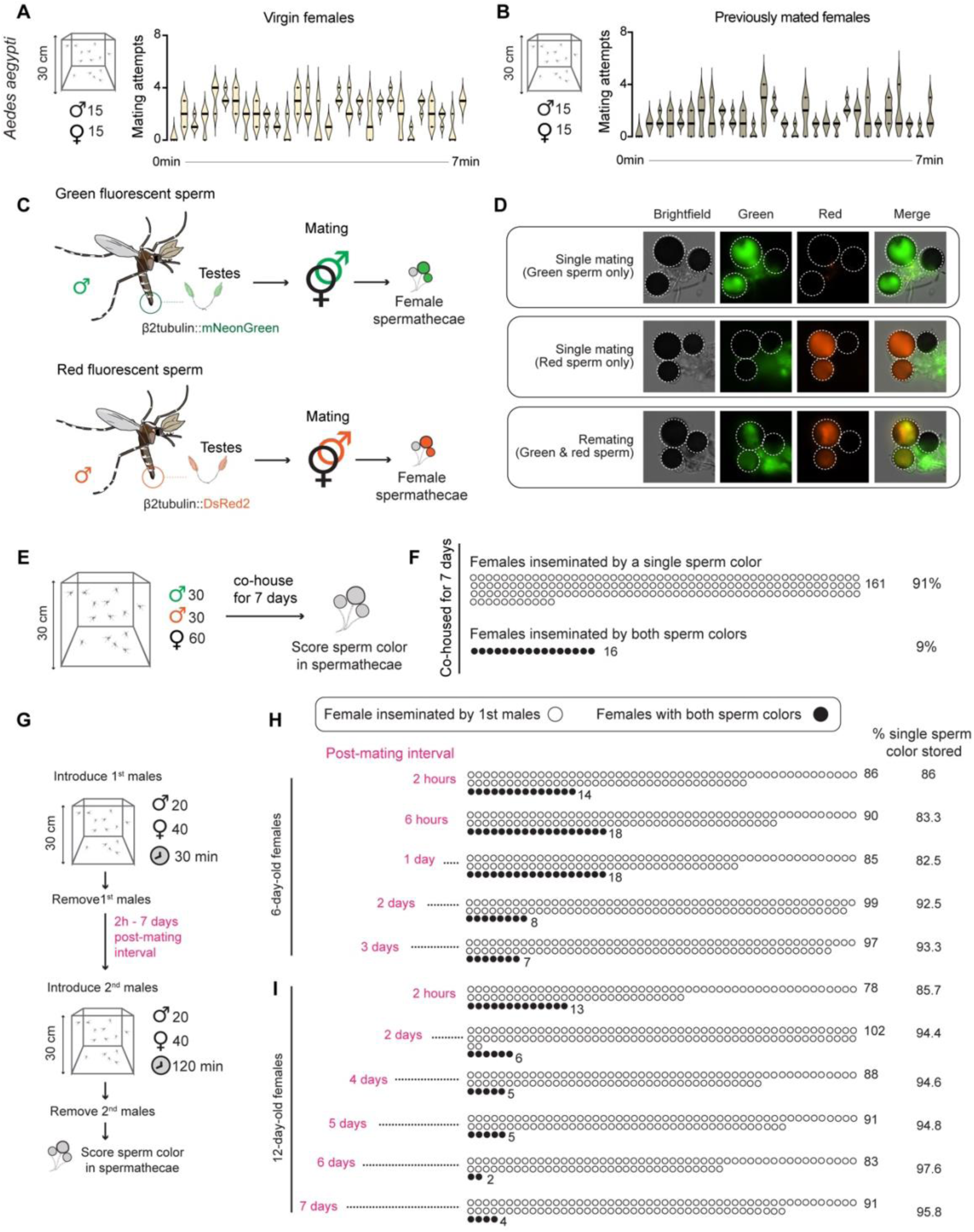
Males persistently attempt to copulate with females, while the female’s remating block is established rapidly. **A. B.** Mating attempts observed for 7 minutes in cages of males and (**A**) virgin or (**B**) previously mated females. N=3 trials, 15 male and 15 female mosquitoes each. Black line indicates the median point. **C.** Schematic of transgenic fluorescent sperm lines expressing mNeonGreen or dsRed2 fluorescent proteins. **D.** Example images of female spermathecae containing green-fluorescent sperm (top), red-fluorescent sperm (middle), or both (bottom). Note green autofluorescence of spermathecal ducts in all samples. **E.** Schematic of experimental setup for testing remating rates following 7-days of co-housing with males with two different fluorescent sperm colors. **F.** Scoring of single or double sperm color of spermathecae from 7-day co-housed females. Each circle is one female. N=3 trials with 60 females each. **G.** Schematic of experimental setup for testing remating rates following introduction of males with two different fluorescent sperm colors at various post-mating intervals. **H. I**. Scoring of single or double sperm color in spermathecae from (**H**) 6-day-old and (**I**) 12-day old females at different post-mating intervals. Data are presented as total scored females across the 3 trials of the experiment. Each circle is one female. N=3 trials with 40 females each. See also **Figure S1**.

*Aedes aegypti* female mosquitoes have three spermathecae, only two of which are typically filled with sperm^7,8^. When we mated transgenic males to wild-type females, we observed rare instances of both red fluorescent and green fluorescent sperm in the spermathecae, which we define as evidence of remating (**Figure 1D**). 91% of females continually co-housed with a mixed group of males for 7 days had a single sperm color in their spermathecae (**Figure 1E,F**), confirming prior observations that most female *Aedes aegypti* females mate only once in a lifetime^3^. A limitation of this experiment is the inability to distinguish single inseminations from multiple inseminations with sperm of the same color, potentially leading to underestimation of the actual remating rates. To address this, we performed experiments in which females were sequentially exposed to males with different sperm colors. We removed the first males and replaced with second males of a different sperm color at different post-mating intervals (**Figure 1G**). We found that 86% of females have an established remating block two hours after mating, and this increased to 95.8% at 7 days post mating (**Figure 1H,I**). The remating block was similar in younger and older females (**Figure 1H,I**). Our results show that most female *Aedes aegypti* block remating rapidly after a single mating event, with a small number of females capable of mating more than once.

### Female genital behavior controls mating in *Aedes aegypti*

What is the behavioral basis of the remating block? Video recordings of 3-dimensional free flight interactions (**Figure S2A-I** and **Extended data Video 1**^29^) and contact interaction of a string-tethered female and a free flying male (**Figure 2A-H**, **Figure S2J-L** and **Extended data Video 1**^29^) revealed that previously mated females spent less time in contact with males when in free flight (**Figure S2B-D**), and that previously mated females kick the male more (**Figure 2F**). Although the male contacted the genitalia of the female numerous times regardless of her mating status (**Figure 2E**), in most of the cases these genital contact events remained superficial. A tighter genital connection was only observed in virgins^4^, which we term *genital interlocking*. Genital interlocking was observed only once per virgin female and was never observed in previously mated females (**Figure 2G,H**). These observations point to genital interlocking as the critical difference between receptive and unreceptive females, prompting us to investigate how this process is controlled.

**Figure 2:**
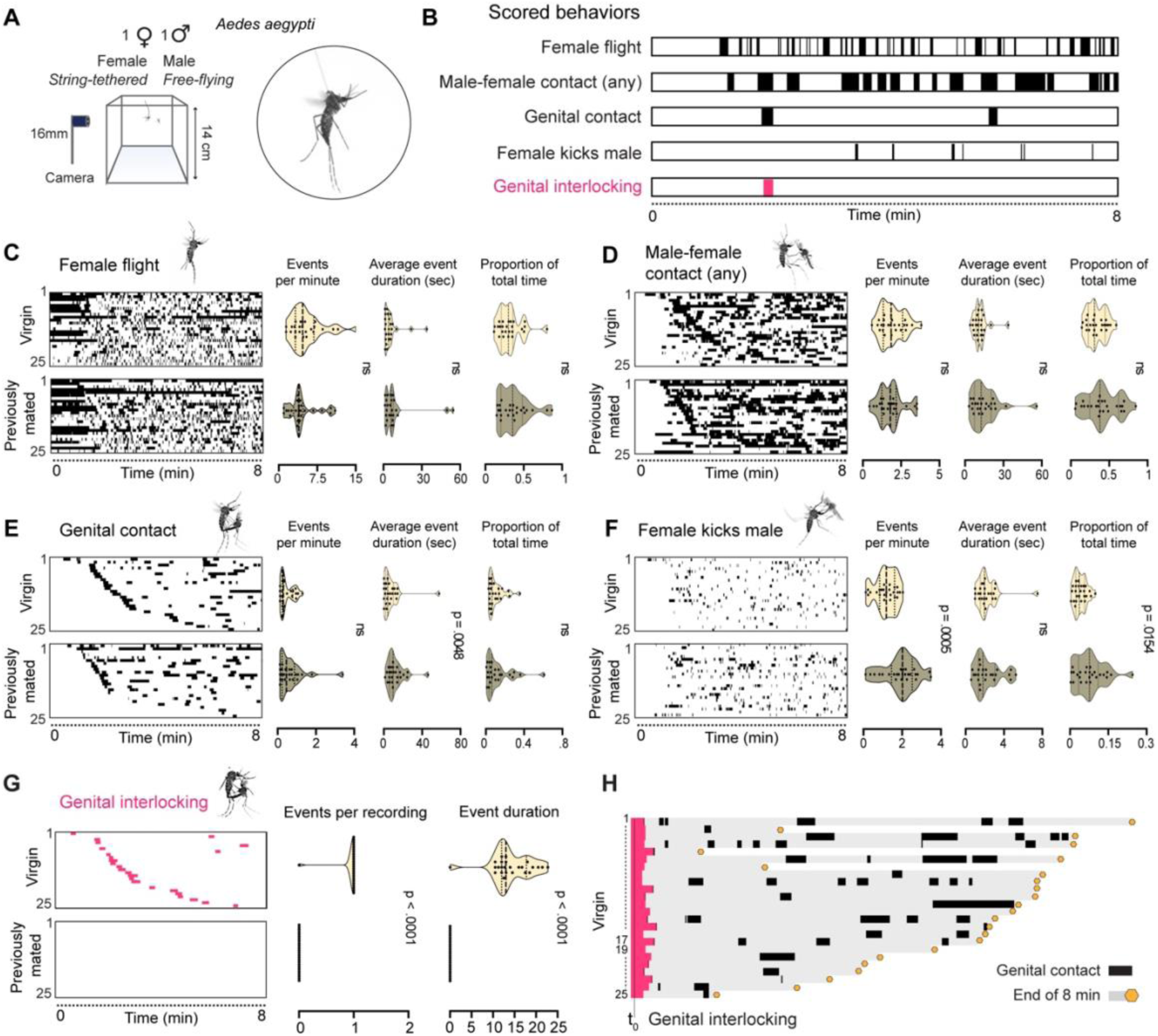
Mating interactions of virgin or previously mated females (string-tethered) and males (free-flying). **A.** Experimental setup and a video still image of a string-tethered female recording, paired with a free flying male. N=25 recordings per condition. **B.** Behavioral categories scored in this assay. Ethogram represents data from virgin recording #8. Note that these data are included in the ethograms in panels i-n below. **C-G**. Heatmaps and violin plots of each scored behavior and a summary of events per minute, average event duration, and proportion of total time per behavior in recordings of virgin and previously mated female. Median (thick dotted line) and 25/75 quartiles (thin dotted line) are indicated. Heatmap is sorted by time until first genital contact event. Each row in the heatmap represents an individual recording, repeated per behavior: (**C**) female flight, (**D**) male-female contact, (**E**) genital contact, (**F**) female kicks male, and (**G**) genital interlocking. Median (thick dotted line) and 25/75 quartiles (thin dotted line) are indicated. Data were analyzed using the Mann-Whitney test. ns=not significant. **H**. Virgin female recordings aligned to the point of genital interlocking and showing ensuing genital contact events until the end of each recording (yellow hexagon). Recording of virgin female #18, that did not interlock during the recording, is excluded from the plot. See also **Figure S2**.

The female vaginal lips are hidden between the ventral side of the 8^th^ sternite and the post-genital plate that sits above the cercal plates (**Figure 3A** and **Figure S3A,B**). The penis-equivalent and sperm transfer organ of the male, the *aedeagus*, is located superior to the secondary genital structures and is sheathed by the *paraproct*^11,30^ (**Figure 3B** and **Figure S3C,D**). For intromission, the aedeagus emerges and fits into the female vaginal tract^11^.

**Figure 3:**
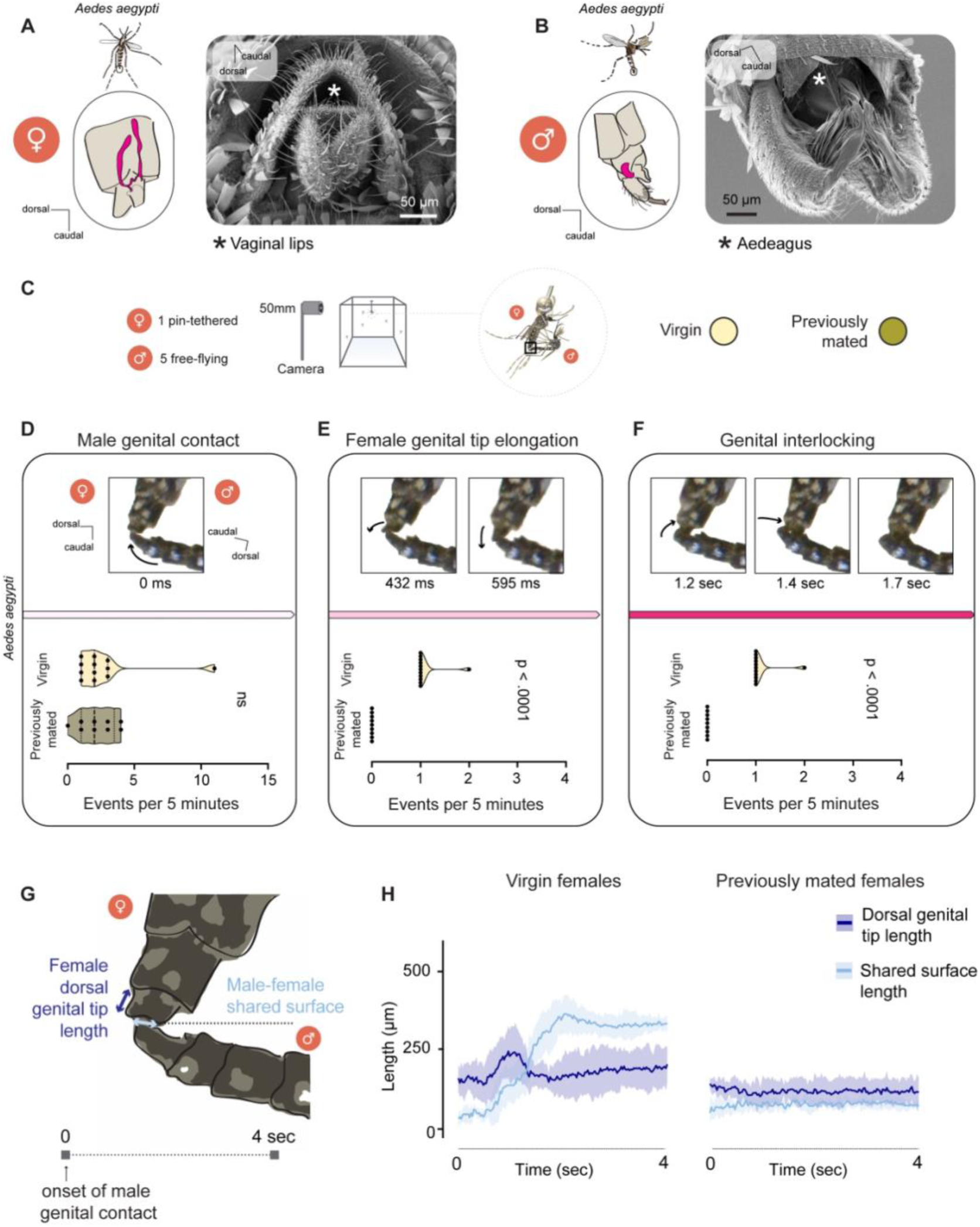
Female genital tip elongation controls mating in *Aedes aegypti* mosquitoes. (**A,B**). Anatomy of the (**A**) female and (**B**) male genital tips represented by schematics (left) and scanning electron microscopy images (right). Magenta structures in schematics indicate (**A**) the female vaginal tract and bursa (sealed by the vaginal lips at the bottom) and (**B**) the male sperm transfer organ, the aedeagus. White asterisks indicate the location of (**A**) the female vaginal lips and (**B**) the male aedeagus, both of which are covered by other genital structures. (**C**). Experimental setup for recording genital interactions of a pin-tethered female and free-flying males. (**D-F**). Representative video still images of the tips of the male and female during the three major steps towards successful mating in *Aedes aegypti* mosquitoes and number of events per 5 minutes of video recordings of (**D**) male genital contact, (**E**) female genital tip elongation, and (**F**) genital interlocking in virgin and previously mated female video recordings. n=12 virgin and n=9 previously mated video recordings with 1 female and 5 males. Time stamp represents time since male genital contact. Median (thick dotted black line) and 25/75 quartiles (thin dotted black line) are indicated. ns=not significant (Mann-Whitney test) **(G)**. Schematic showing quantification strategy for measuring genital behavior. (**H**). Quantification of indicated genital behaviors in virgin and previously mated female video recordings. n=6 tracked video recordings per condition. Average traces are shown with 95% confidence intervals. See also **Figure S3**.

Prior studies of mosquito copulation described a multi-step process for the full formation of a genital connection and intromission that was presumed to be entirely guided by the male^11,31,32^. Similar to our results, prior work showed that mating attempts between males and previously mated females are limited to a superficial genital connection^4^. To better understand the process that leads to successful mating and genital interlocking, we video recorded male-female interactions in high resolution, specifically focusing on the genital behavior of the male and the female. This was accomplished by tethering the female to a metal pin to limit her movement and provide a consistent focal point for our video recordings (**Figure 3C** and **Extended data Video 1**^29^). We recorded videos of virgin and previously mated females interacting with free-flying males and scrutinized these video recordings frame-by-frame for male-female genital interactions. With this approach, we discovered three major steps that lead to successful mating in *Aedes aegypti* mosquitoes. First, the male genitalia contact the female genital tip (**Figure 3D**). Second, in response to genital contact by the male, virgin females elongated their genital tip (**Figure 3E**), a previously undescribed female mosquito genital behavior. Third, this act of female genital response was followed by successful genital interlocking. Importantly, previously mated females never elongated their genital tips in response to male genital contact and never engaged in genital interlocking (**Figure 3F**).

To quantify these genital movements, we developed a manual tracking strategy for measuring genital behavior in the four seconds after male genital contact was initiated. We extracted measurements of the female dorsal genital tip length as a readout of female genital tip elongation length, and the shared surface between male and female genitalia as a readout of genital interlocking (**Figure 3G-H**).

We found that the female elongated her genital tip to approximately twice its resting length, reaching an average maximum length of 280 µm. Female genital tip elongation always preceded the genital interlocking-associated increase in the connection surface between the male and the female, reaching an average of 390 µm of connection during genital interlocking (**Figure 3H** and **Figure S3E,F**). These dynamics were only observed in genital interactions of virgin females with males. Previously mated females showed no change in either female dorsal genital tip length or male-female shared surface (**Figure 3H** and **Extended data Video 1**^29^). Video recordings of sequential interactions of tethered females with male groups with different fluorescent sperm colors revealed that two females that elongated their genital tip twice were inseminated by sperm of both colors (**Figure 3G-K**). Together, these observations strongly suggest that female mosquitoes actively control mating through elongation of their genital tip.

### Secondary male genital structures trigger the female genital response

Having shown that female genital tip elongation can follow male genital contact, we asked what about this male contact triggers the female to elongate her genital tip. To answer this, we analyzed male-female genital interactions prior to the female genital response (**Figure 4A**). We observed that male genital contact is characterized by the male inserting hook-like genital structures known as the gonostyli (singular: gonostylus) into the female genital tip (**Figure 4A,B**). The gonostyli are secondary male genital structures that are not directly involved in intromission and insemination and rest on the sides of the female cercal plates during copulation^11,32^ (**Figure 4C**). Frame-by-frame examination of videos of pin-tethered females and free-flying males showed that when the male was about to initiate male genital contact, the gonostyli extended, slid into the ventral side of the female genital tip, and vibrated (**Figure 4D** and **Extended data Video 1**^29^). In virgin females, this sequence was followed by female genital tip elongation. This response was rapid, with females beginning their tip elongation an average of 784 milliseconds after initiation of male genital contact. In previously mated females, males vibrated their gonostyli against the female genital tip for the duration of male genital contact, but the females did not respond (see **Extended data Video 1**^29^). The gonostyli are rapidly evolving structures that assume different shapes across the mosquito tree of life^36^ (**Figure 4E**)

**Figure 4:**
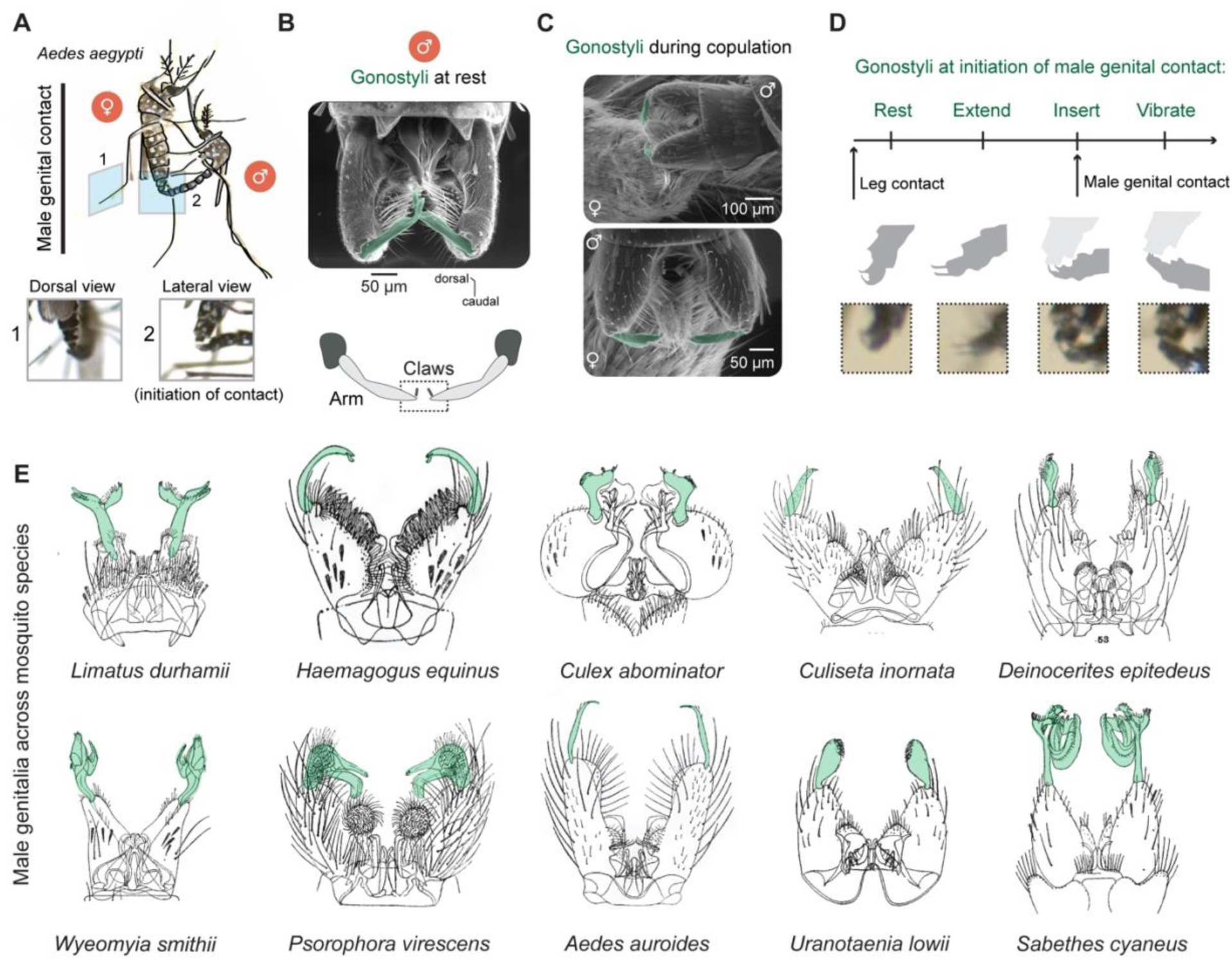
Gonostyli in mating interactions and across species. **(A)**. Schematics and photographs of dorsal and lateral views of male genital contact. **(B)**. Scanning electron microscopy image of the male genitalia, with gonostyli pseudo-colored in green (top) and a schematic of the gonostyli (bottom). **(C)**. Cryo-scanning electron microscopy images of mosquito genitalia during copulation. Gonostyli are pseudo-colored green. **(D)**. The position and behavior of the gonostyli prior to and during male genital contact. (**E**) Scientific drawings of male genitalia of the indicated species across *Culicine* genera. Putative gonostyli structures colored in green. (reproduced from Howard et al. 1912)^36^

To determine the role of gonostyli insertion prior to female genital tip elongation we tested the effect of surgical gonostyli manipulation on male-female genital interactions with a pin-tethered female (**Figure 5A** and **Figure S4A,B**). Importantly, we observed that gonostyli-manipulated males recovered well from the procedure and attempted to copulate with females to similar or higher degrees compared to “mock” treated males (**Figure S4C**). In all 14 video recordings, virgin females paired with mock-treated males elongated their genital tips in response to male genital contact and genital interlocking was achieved in all cases (**Figure 5B**). In contrast, in 11 of 14 video recordings of virgin females paired with males with gonostyli removed, the females did not elongate their genital tips in response to male genital contact and genital interlocking was never achieved (**Figure 5C**). In the three cases that females elongated their genital tips in response to male genital contact with gonostyli-less males, the male attempted to genitally interlock but failed. Of 14 video recordings of virgin females paired with males with dislocated gonostyli, none elongated their genital tip in response to male genital contact and genital interlocking was never attempted or achieved (**Figure 5D**).

**Figure 5:**
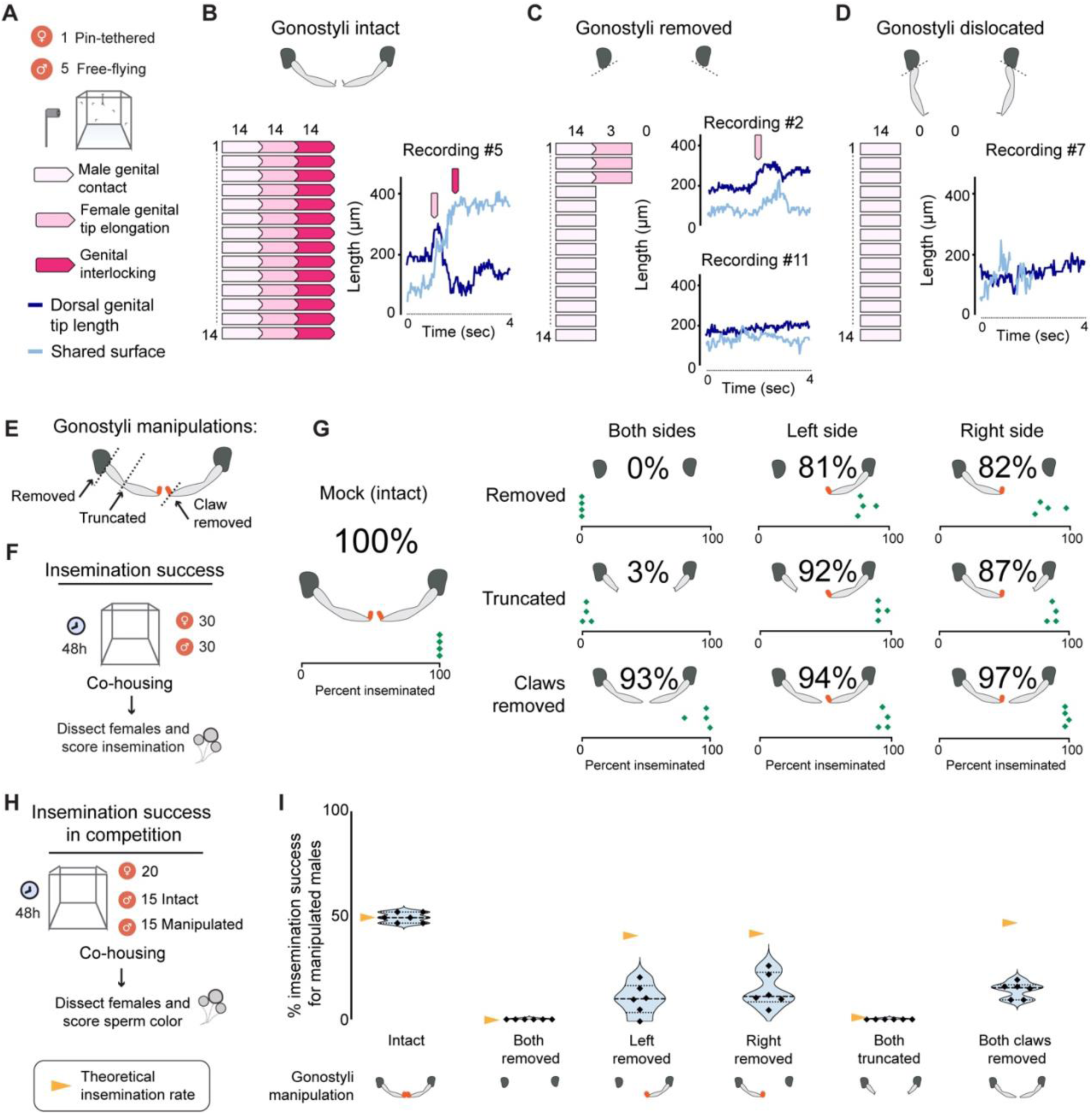
Gonostyli trigger female genital tip elongation and are required for successful insemination. (**A**). A schematic of the experimental setup for testing the effect of gonostyli manipulation on genital behavior (top) and the indications of the different genital interaction steps and measurements (bottom). (**B-D**). Genital behavior and representative tracking dynamics in mating interactions with (**B**) mock-treated males or male with (**C**) gonostyli removed or (**D**) dislocated (motionless). N=14 5-minute video recordings per condition. (**E**). Schematic of gonostyli manipulations examined for insemination success. (**F**). A schematic of the experimental setup for testing insemination success of mock treated and manipulated gonostyli males. (**G**). Insemination rates of females co-housed with mock treated and manipulated gonostyli males. Overall averaged percent insemination success across trials and dot plots showing insemination success rates per trial are displayed. N=4 trials per condition. (**H**). A schematic of the experimental setup for testing insemination success in competition. (**I**). Insemination success rates of intact (mock-treated) males or males with manipulated gonostyli when challenged by competition with mock-treated males. Median (thick dotted black line) and 25/75 quartiles (thin dotted black line) are indicated. Yellow triangle indicates theoretical insemination success rates calculated from a no-competition scenario (panel **G**). N=3 trials with each condition tested twice per trial, switching between green and red sperm males as the manipulated group. See also **Figure S4**.

Gonostyli are bilaterally symmetric curved cuticular structures with two arms and one small claw at the tip of each arm. Using a free-flight setup, we tested how finer gonostyli manipulations affect insemination success (**Figure 5E-G** and **Figure S4B-C**). Across replicates, we observed that mock-treated males successfully inseminated 100% of the females they were paired with. Bilateral removal or truncation of both sides of the gonostyli nearly eliminated insemination success, whereas removal of only the two claws had a modest impact on insemination success (**Figure 5G**). Unilateral removal of a gonostylus, truncation, or removal of a claw reduced insemination success 3-19% relative to mock-treated males, indicating that for most females, one fully intact gonostylus is sufficient to induce female genital tip elongation (**Figure 5G**).

To ask if gonostyli manipulation impacts insemination success in the context of competition with males with intact gonostyli, we paired females with mock-treated males with one sperm color and surgically manipulated males with a second sperm color (**Figure 5H**). Unsurprisingly, males with bilateral gonostyli removal or truncation always lost to mock-treated males. Remarkably, males with unilateral gonostylus removal who were ∼80% successful in inseminating females without competition saw their insemination success rate drop to an average of 11-15% in competition compared to the theoretical insemination success rate of 40% calculated from a no-competition scenario (**Figure 5I**). Even more dramatically, males lacking only the claws at the tip of their gonostyli who were 93% successful in inseminating females in a non-competitive environment were outcompeted by mock-treated males and had an average insemination success rate of 15% compared to the theoretical no-competition success rate of 46.5% (**Figure 5I**). Together, these results suggest a critical role for the gonostyli in triggering female receptivity via the female genital tip elongation response.

### Initiation of successful mating in *Aedes albopictus* mosquitoes

The system of a male genital structure that promotes female receptivity and successful insemination suggests a “lock and key” mechanism between male and female genitalia. In this framework, rapid evolution of animal genitalia can safeguard the species barrier through structural compatibility and incompatibility^33–35^. This variation may be particularly important to prevent cross-species mating in species that share the same habitat.

We asked how genital interactions during initiation of successful mating vary in another *Aedes* species, *Aedes albopictus*, which has expanded globally over the past 50 years and now frequently shares territories with *Aedes aegypti*^37^. While the structure of the female genital tip is similar between the two species (**Figure S5A**), male *Aedes albopictus* gonostyli are significantly larger in both length and thickness compared to male *Aedes aegypti* gonostyli (**Figure 6A** and **Figure S5B**).

**Figure 6:**
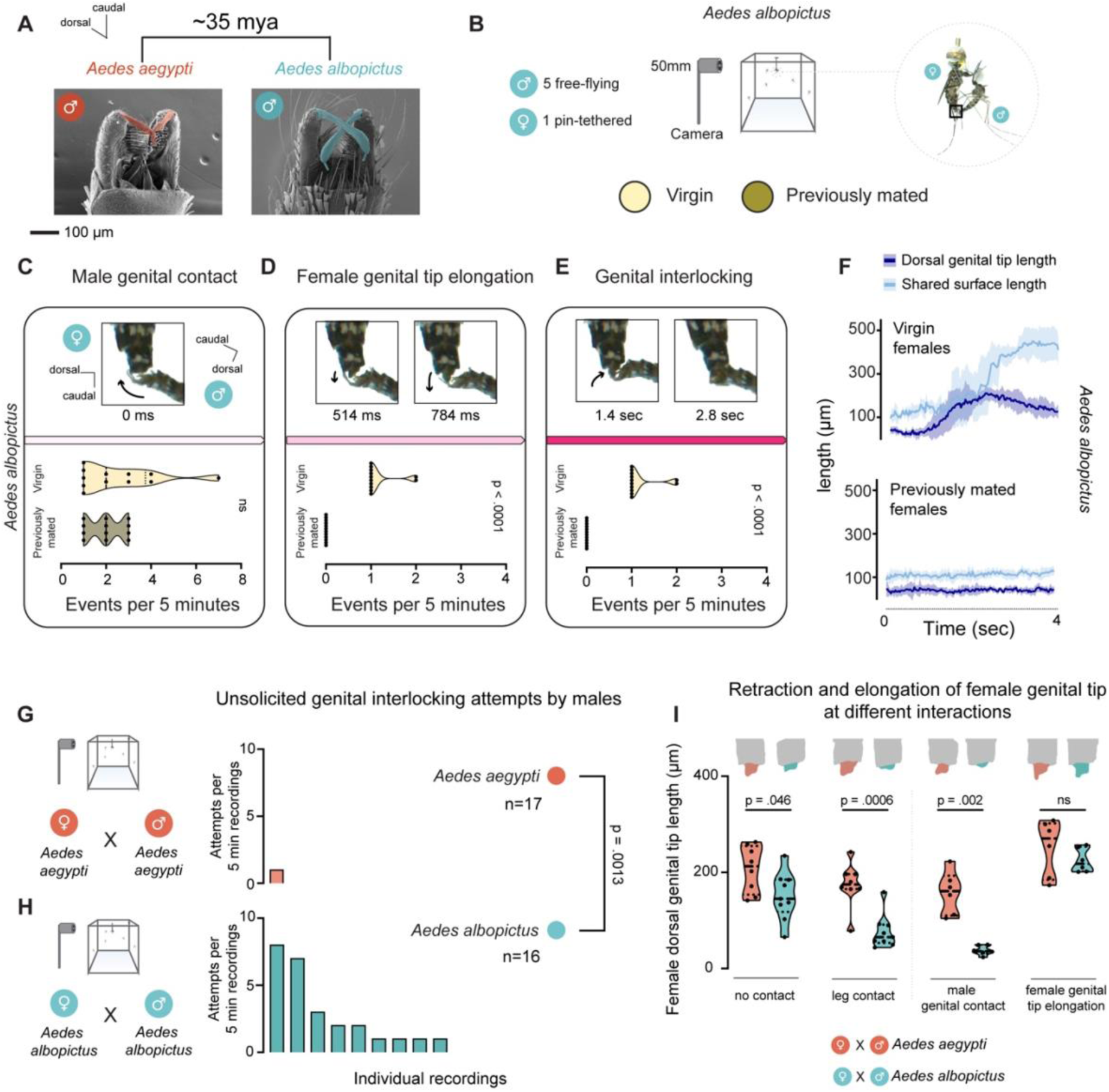
Comparison of genital interactions and behaviors in *Aedes albopictus* and *Aedes aegypti*. (**A**). Scanning electron microscopy images of the ventral side of *Aedes aegypti* and *Aedes albopictus* male genitalia. Gonostyli are pseudo-colored orange (*Aedes aegypti*) and turquoise (*Aedes albopictus*) (**B**). Experimental setup for recording of genital behavior of a pin-tethered female and free-flying males. (**C-E**). Representative video still image of the tips of the male and female during the three major steps towards successful mating in *Aedes albopictus* mosquitoes and number of events per 5 minutes of video recordings of (**C**) male genital contact, (**D**) female genital tip elongation, and (**E**) genital interlocking – in virgin and previously mated female video recordings. N=12 video recordings with 1 female and 5 males per condition. Time stamp represents time since the depicted video still image of male genital contact. Median (thick dotted black line) and 25/75 quartiles (thin dotted black line) are indicated. Data were analyzed using the Mann-Whitney test. ns=not significant. (**F**). Average quantification and 95% confidence interval of female dorsal genital tip length and male-female shared surface length in genital interactions between males and virgin or previously mated female video recordings. n=6 video tracked per condition. (**G,H)**. Unsolicited genital interlocking attempts by males in same-species mating in (**G**) *Aedes aegypti* and (**H**) *Aedes albopictus* mosquitoes. Each bar represents the number of events per examined recording. Data are pooled from video recordings of virgin and previously mated females. Data were analyzed using the Mann-Whitney test. ns=not significant. (**I**). Female dorsal genital tip length in *Aedes aegypti* and *Aedes albopictus* at different stages of interactions with males. N=8,9 analyzed pre- and post-leg contact video recordings in *Aedes aegypti* and *Aedes albopictus* (respectively). N=6 analyzed video recordings in male genital contact and female genital tip elongation. Median (thick dotted black line) and 25/75 quartiles (thin dotted black line) are indicated. Data were analyzed using the Mann-Whitney test. ns=not significant. See also **Figure S5**.

We examined male-female genital interactions in *Aedes albopictus* using our high-resolution video recording setup of a pin-tethered female and free-flying males (**Figure 6B**). We found that *Aedes albopictus* mating follows the same behavioral sequence of male genital contact, female genital tip elongation, and genital interlocking that we observed in *Aedes aegypti* (**Figure 6C-E** and **Extended data Video 1**^29^). In *Aedes albopictus*, female genital tip elongation is required for genital interlocking and is absent in previously mated females, exactly as we observed in *Aedes aegypti* (**Figure 6D,F**, **Figure S5C,D** and **Extended data Video 1**^29^). This demonstrates that female control over mating, and the behavioral pathway leading to successful mating, is shared in mosquitoes with ∼35 million years of evolutionary distance.

Despite these similarities, we identified three fundamental differences in genital behaviors of the two species. First, during male genital contact, the large gonostyli of *Aedes albopictus* males were positioned dorsally and externally (**Figure 6C**) while the smaller *Aedes aegypti* gonostyli were positioned ventrally and internally to the female genital tip (**Figure 3D** and **4D**). Second, *Aedes albopictus* males tried to genitally interlock with females even in the absence of female genital tip elongation, a behavior that we observed only once in *Aedes aegypti* (**Figure 6G,H** and **Extended data Video 1**^29^). Third, while elongation of the female genital tip reached similar lengths in both species during the female genital response, only *Aedes albopictus* females displayed dramatic retraction of their genital tip at all steps preceding the genital tip elongation response that signals their receptivity (**Figure 6I**).

### Lock and key functionality in cross-species genital interactions

Having shown that genital interactions of male and female mosquitoes differ between *Aedes aegypti* and *Aedes albopictus* during initiation of successful mating, we asked how these differences translate to genital interactions between the two species. Because these species mate at low frequency under laboratory conditions, we examined genital interactions using pin-tethered pairs to manually bring the male into close contact with the female (**Figure 7A**).

**Figure 7:**
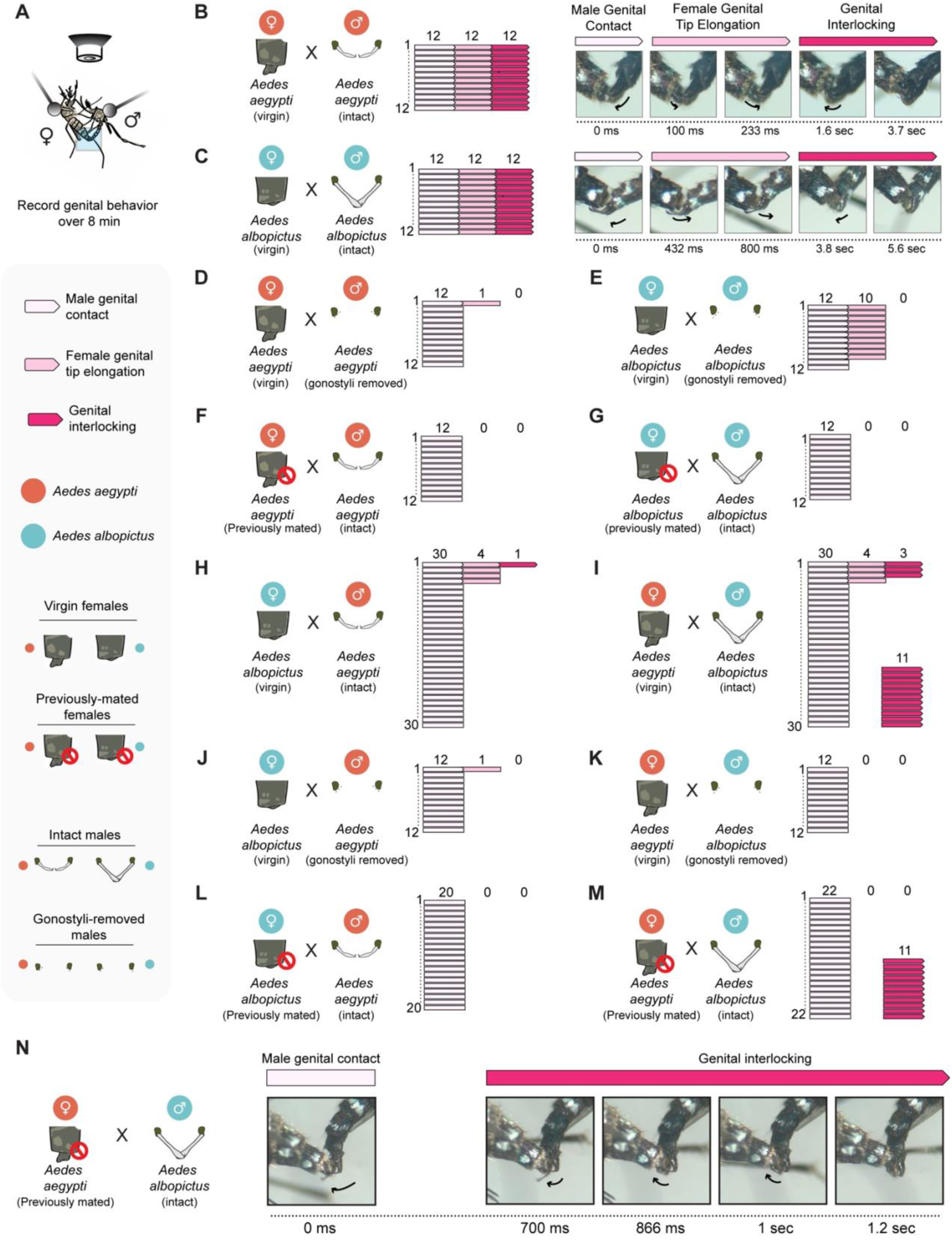
Same-species and cross-species genital interactions in *Aedes aegypti* and *Aedes albopictus*. (**A**). A schematic of the experimental setup and legend for pin-tethered male and female genital interaction video recordings. (**B-G**). Same-species genital interactions of (**B**,**C**) virgin females and intact males together with video still images of the genital interactions, of (**D**,**E**) virgin females and males with their gonostyli removed, and of (**F**,**G**) previously mated females and intact males. (**H-M**). Cross-species genital interactions of (**H**,**I**) virgin females and intact males together with video still images of the genital interactions, of (**J**,**K**) virgin females and males with their gonostyli removed, and of (**L**,**M**) previously mated females and intact males. (**N**). Video still images of genital interactions between previously mated *Aedes aegypti* female and intact *Aedes albopictus* male. Female genital tip elongation response is bypassed directly to genital interlocking. The number of video recordings per condition is indicated on each panel.

Same-species genital interactions of females and intact males followed the same dynamics that we observed before. In both species, the animals followed the behavioral sequence of male genital contact, female genital tip elongation, and genital interlocking (**Figure 7B,C** and **Extended data Video 1**^29^). While the gonostyli trigger female genital tip elongation in *Aedes aegypti* (**Figure 5A-D** and **Figure 7D**), *Aedes albopictus* males without gonostyli were able to successfully induce female genital tip elongation in 10 of 12 pairs but were unable to proceed to genital interlocking (**Figure 7E**). This suggests that *Aedes albopictus* males use other unknown mechanisms to trigger female genital tip elongation in females of their own species. As observed before, previously mated females of both species never elongated their genital tip, and the pair did not proceed to genital interlocking (**Figure 7F,G**).

In cross-species mating, when intact *Aedes aegypti* males with their small gonostyli were paired with virgin *Aedes albopictus* females with their retracted genital tip, females rarely elongated their genital tip (4 of 30 pairs) and genital interlocking was achieved in only one of these 4 pairs (**Figure 7H**).

When *Aedes albopictus* males with their big gonostyli were paired with *Aedes aegypti* females with their extended genital tip, female genital tip elongation was a similarly rare event (4 of 30 pairs) although successful genital interlocking occurred in 3 of these pairs (**Figure 7I**). Strikingly, in 11 of 30 pairs, *Aedes albopictus* males bypassed *Aedes aegypti* female genital tip elongation and moved directly to genital interlocking (**Figure 7I**). This unique ability of male *Aedes albopictus* required intact gonostyli (**Figure 7J,K**).

Given that *Aedes albopictus* are able to bypass the female genital response, we reasoned that these males should be able to overcome the remating block of previously mated *Aedes aegypti* females. We therefore carried out cross-species mating experiments with previously mated females. We first found that *Aedes aegypti* males were unable to trigger female genital tip elongation or proceed to genital interlocking with previously mated *Aedes albopictus* females. We conclude that *Aedes aegypti* males are unable to remate with previously mated females from their own species or the other species (**Figure 7F,L**). In contrast, we found that *Aedes albopictus* males were able proceed to genital interlocking with previously mated *Aedes aegypti* females in half (11 of 22) of the interacting pairs (**Figure 7M,N** and **Extended data Video 1**^29^). In all cases, *Aedes albopictus* males bypassed female *Aedes aegypti* genital tip elongation and proceeded immediately to cross-species genital interlocking with these previously mated females. Male *Aedes albopictus* were never observed to bypass the female genital response when attempting to mate with females of their own species (Figure 6C-F, Figure 7G, **and** Figure S5C,D).

Together, these results reveal a notable asymmetry in cross-species mating: *Aedes albopictus* males can bypass the female response and break the remating block across species, potentially through mechanical advantage conferred by their larger gonostyli and the extended genital tip of female *Aedes aegypti*.

## Discussion

In this study we established that a female genital behavior controls mating in *Aedes aegypti* and *Aedes albopictus* mosquitoes. In *Aedes aegypti*, the female genital response is triggered by the gonostyli, rapidly evolving secondary male genital structures. We identified key differences in genital interactions of *Aedes aegypti* and *Aedes albopictus* and found that *Aedes albopictus* males, with their larger gonostyli, can bypass the genital response of *Aedes aegypti* females but not females of their own species.

It is not known how the interaction of the male gonostyli with the female genital tip unlocks the female genital response. Our observations support several possible mechanisms that are not mutually exclusive. First, the impressive variability and rapid evolution of the gonostyli structure across species^36,38^ suggest that information might be relayed by the shape of the gonostylus itself. Second, physical interactions of the gonostyli with female genitalia could provide an additional component to the code. Prior to female genital tip elongation, the gonostyli are inserted into the female genital tip and vibrate rapidly. It is possible that the vibration rhythm of the gonostyli is recognized by the female via a yet unknown mechanosensory pathway and creates a unique code across species. Third, it is also possible that gonostyli transmit contact chemosensory cues that together with its vibration pattern could trigger the female genital response.

It will be interesting to understand how the female senses male gonostyli stimulation and the regulation of the motor response governing genital tip elongation. The cercal and the post-genital plates at the tip of the female genitalia form a triangular structure that is densely covered with sensilla. These sensilla are described to be mostly mechanosensory^39^. The post-genital plate is typically pushed by the gonostyli during male genital contact and is in contact with them during the vibration phase. It is possible that genital plate sensilla have a role in recognizing gonostyli insertion and vibration, or they may recognize additional cues including chemical substances. Alternatively, and perhaps synergistically, the tips of the gonostyli contact a membranal-like tissue at the base of the post-genital plate and near the vaginal lips. This tissue could be internally innervated to sense and recognize gonostyli stimulation. Understanding the sensory components of the circuitry that control female genital tip elongation would provide better understanding of the interactions between male gonostyli and the female genital tip, as well as access to neuronal regulation of mating in female mosquitoes, and how this regulation changes rapidly after a single mating event.

Our findings align with the “lock and key” theory for genital evolution. This framework, first formulated in 1844^33,34^, states that the rapid evolution of animal genitalia is a mechanism for safeguarding the species barrier through structural compatibility and incompatibility. In our system, the gonostyli function as the key to unlock the female genital response in *Aedes aegypti* to initiate successful mating. While the male shows morphological variation of the gonostyli across the two species studied here, the female side of the mechanism appears to be mostly behavioral. This system better matches updated versions of the “lock and key” mechanism that focus on female sensory reception and behavior over structural differences^35^. Understanding the interactions or structures that serve to stimulate the female genital responses in other systems, and how the rapid evolution of the male genitalia corresponds to the female sensory capacity and behavioral response across species, would provide fascinating insights into the evolution of mating systems, male-female co-evolution, and cross-species competition.

Over the past 50 years, *Aedes albopictus* mosquitoes have successfully invaded Europe and the Americas from their original geographic range in Asia. They readily adapt to new environments and outcompete other mosquito species in shared territories^18,40^. Our finding that male *Aedes albopictus* can bypass the female genital response in *Aedes aegypti* may provide a mechanism for the competitive and invasive capabilities of *Aedes albopictus.* Cross-species mating between *Aedes aegypti* and *Aedes albopictus* does not yield viable progeny and prevents *Aedes aegypti* females from mating again^23,24^. While the discovered lock-and-key mechanism functions in *Aedes aegypti* male-female genital interactions, our findings reveal that *Aedes albopictus* males can coercively bypass this system when encountering *Aedes aegypti* females. This represents interspecific sexual coercion that overrides, rather than replaces, the normal lock-and-key system. Future investigation is needed to understand what additional cues might play a role in these cross-species interactions and the female’s ability to elongate her genital tip specifically in response to conspecific males. Finally, in some regions of the world *Aedes albopictus* are weak “satyrs”^41^. It would interesting whether variation in gonostyli shape and behavior might underlie this difference.

*Aedes aegypti* and *Aedes albopictus* mosquitoes transmit dangerous arboviruses such as dengue, Zika, and chikungunya to humans^42,43^. Reducing mosquito population sizes is a major public health priority to lower the burden of disease. Much of the effort to control mosquito population relies on mating as a point of intervention. The Sterile Insect Technique and Incompatible Insect Technique involve releasing millions of sterile or Wolbachia-infected males to targeted areas^44^. When these males successfully mate with wild females, the females produce no viable progeny and are effectively sterilized^45^, reducing mosquito population size and viral transmission. In this study we describe the process leading to successful mating in *Aedes aegypti* and *Aedes albopictus* mosquitoes and the role of the female in controlling it. Our findings provide a framework for enhancing mosquito control methods by accounting for the newly discovered role of the female in mating and the male’s ability to induce this receptivity.

## Resource Availability

### Lead contact

Requests for further information and resources should be directed to and will be fulfilled by the lead contact, Leah Houri-Zeevi.

### Materials availability

Plasmids generated in this study have been deposited to Addgene (#236085, #236086). All mosquito strains generated in this study are freely available on request.

### Data and Code Availability

- Data are publicly available at the paper’s GitHub depository^46^ and Zenodo record #15133603^29^.
- All original code has been deposited at the paper’s GitHub depository^46^ and is publicly available as of the date of publication.
- Any additional information required to reanalyze the data reported in this work is available from the Lead Contact upon request.

## Acknowledgements

We thank members of the Vosshall Lab and Philipp Brand, Tomas Kay, Maximilian M.L. Knott, Orli Snir, and Itai A. Toker for discussion and comments on the manuscript; Gloria Gordon, Libby Mejia, and Melissa Dallesandro for strain maintenance, and Mona Liu for early contributions to the project. We thank Peter Atkinson, Rob Harrell, and Laura Harrington for providing the PB{3xP3-EGFP; β2Tubulin-dsRed2} plasmid. We thank the F.M Kirby Foundation for support in acquiring the JEOL JSM-IT500HR scanning electron microscope. This work was supported by a Junior Fellowship of the Simons Society of Fellows and the European Molecular Biology Organization long-term fellowship (EMBO ALTF 1103-2019) (L.H-Z.), and a graduate fellowship from the Price Family Center for the Social Brain and from the Boehringer Ingelheim Fonds (J.R.). L.B.V. is an Investigator of the Howard Hughes Medical Institute.

## Author contributions

L.H-Z. carried out or developed and supervised all the experiments and data analyses in the paper with additional contributions from co-authors. M.M.W. carried out the free flight and pin-tethered video recordings and gonostyli manipulation experiments. J.R. developed the 3D tracking analysis for mosquitoes in free flight and pose estimation analysis for string-tethered video recordings (**Figure S2**). A.S. and H.A.P. processed the mosquito samples for scanning electron microscopy and obtained the images. L.H-Z. and L.B.V. together conceived the study, designed the figures, and wrote the paper with input from all authors.

## Open access statement

This article is subject to HHMI’s Open Access to Publications policy. HHMI lab heads have previously granted a nonexclusive CC BY 4.0 license to the public and a sublicensable license to HHMI in their research articles. Pursuant to those licenses, the author-accepted manuscript of this article can be made freely available under a CC BY 4.0 license immediately upon publication.

## STAR★Methods

### EXPERIMENTAL MODEL AND STUDY PARTICIPANT DETAILS

#### Animals used in this study

The following mosquito species and strains were used in this study:

*Aedes aegypti*: Liverpool wild-type laboratory strain (LVP), PB{3xP3-EGFP; β2Tubulin-dsRed2} (red fluorescent sperm strain, LVP background), and PB{3xP3-dsRed2; β2Tubulin-mNeonGreen} (green-fluorescent sperm strain, LVP background). *Aedes albopictus*: Foshan inbred strain Pavia A (FPA) wild-type laboratory strain. Unless otherwise noted, animals used in this study were between 3 days old and 12 days old, defined as days after adult eclosion. Both males and females were sugar fed and unless otherwise specified females were not blood fed. We define “previously mated” females as those that were at least 24 hours post mating. The LVP strain (*Aedes aegypti* wild type) was initially collected in west Africa and has been maintained at the Liverpool School of Tropical Medicine since 1936^47^. The FPA strain (*Aedes albopictus* wild type) was initially isolated in southeast Asia in 1981 and has been cultivated in the lab since then^48^

### METHOD DETAILS

#### Mosquito rearing

Mosquito strains were reared in a constant temperature and humidity environmental room at 26°C with 70 to 80% relative humidity and daily 14-hour light and 10-hour dark cycle (lights on at 7 a.m.). All behavioral experiments, except pin-tethered male and female mating (**Figure 7**), were performed at 26°C with 70 to 80% relative humidity. Pin-tethered male and female mating experiments were performed at ambient temperature and humidity. Experiments were performed between 9 a.m. and 5 p.m. Embryos were hatched in hatching broth, which was made by crushing one pellet of fish food (TetraMin Tropical Tablets, Pet Mountain, YT16110M) using a mortar and pestle, mixing the powder into 850 ml of deionized water and autoclaving it. The following day, 2500 ml of deionized water were added to the hatched larvae, and they were fed two fish food tablets per day until pupation. Pupae were sexed based on size and shape and male and female pupae placed into separate small cups (VRW, Cat. No. 89009-664) with deionized water. Since even a single male could “contaminate” the virgin female cage, the female pupal cups were covered with a net until eclosion. Upon eclosion, animals were aspirated out in small groups and verified to be female before releasing into designated virgin female cage (30cm^3^ BugDorm-1 Insect Rearing Cage, MegaView Science). After eclosion, adult mosquitoes had access to 10% sucrose (w/v in distilled water), which was delivered in a Boston clear round 60 ml glass bottle (Fisher Scientific, FB02911944) with sugar dispensed through a saturated cotton dental wick (Richmond Dental, 201205). For egg collection, female mosquitoes were blood-fed on mice or sheep’s blood (HemoStat Laboratories, DSB100) mixture with ATP (20 mM ATP in 25 mM aqueous sodium bicarbonate added to the blood to a 2 mM final concentration) and metal blood pucks as previously described^49^. Eggs were dried at 26°C and 80% humidity for three days, then stored at ambient temperature and humidity for up to 4 months.

#### Transgenic strain generation and rearing

Red and green fluorescent sperm lines were generated using PiggyBac transposition-based transgenesis. The PB{3xP3-EGFP; β2Tubulin-dsRed2} plasmid was kindly provided by Robert Harrell at the Insect Transformation Facility, with the permission of Dr. Peter Atkinson^50^ and Dr. Laura Harrington. Transgenic males with this plasmid produce red fluorescent sperm under the control of the β2Tubulin promoter, which confers marker expression in mature sperm. The 3XP3 promoter is used as a marker of transgene insertion and confers marker expression in the eyes. To generate a new strain that produces green fluorescent sperm, the PB{3xP3-EGFP; β2Tubulin-dsRed2} plasmid was modified using the NEBuilder HiFi DNA assembly kit (NEB E5520S) to replace the fluorescent proteins with dsRed2 under the 3xP3 promoter (eye marker), and mNeonGreen. Full plasmid details and sequences are available at AddGene (plasmids #236086 and #236085). Each plasmid was injected at 200 ng/μL together with PiggyBac transposase mRNA (200 ng/μL) to induce transposition-based genomic integration. Injections were carried out in house using the Zeiss SteREO Discovery V.8 microscope (Zeiss, 495007-9880-010) and Sutter Instrument injection system (Sutter Instrument MPS-200 controller, FG-MPC385; XenoWorks Digital Microinjector Sutter Instrument, FG-BRE) into mosquito embryos collected 1 hour after being laid using a standard injection protocol^51^. Injected G0 male and female animals were reared until adulthood and crossed to LVP animals. G1 animals were screened for the fluorescent eye marker and two founder animals per injected plasmid were isolated. Lines were then backcrossed to LVP for 3 generations before being inbred. After validating that all lines show wild-type levels of mating success and male fertility, one line per injected plasmid was randomly selected for further study. Transgenic animals were sorted as pupae under a fluorescent microscope (Nikon SMZ18, NIS-Elements AR software) at 475 nm (green fluorescence) or 549 nm (red fluorescence) to confirm 3XP3-derived fluorescence from the transgenic insertion in eyes (both sexes) and testes (males) and separated based on sex as described above.

#### Testing transgenic animals (Figure S1)

Reproductive behavior of transgenic males was assessed as follows:

*Insemination success*: per tested group, 50 wild-type females were separately introduced to 50 males of the three different genotypes (wild type, red fluorescent sperm, green-fluorescent sperm) for one hour. Females were cold anesthetized for five minutes and dissected to score sperm presence in their spermathecae. Experiments were carried out in 5 independent biological replicates.

*<u>Egg laying and hatch rate</u>*: per tested group, 60 wild-type females were introduced to 60 males of the different genotypes (wild type, red-fluorescent sperm, green-fluorescent sperm) for a period of two hours. Following the mating period, 10 females of each group were dissected to ensure successful mating. 2 days post mating, the females were blood fed using Sheep’s blood (HemoStat Laboratories, DSB100) mixture with ATP and metal blood pucks as described before^49^. The few females that were not fully engorged were discarded. Fully engorged females were placed in individual vials for egg laying at 5 days after blood feeding and were allowed to lay eggs for 24 hours. Egg-laying vials were made using plastic Drosophila vials (VWR, 25 mm diameter, 95 mm length, 75813–164) with 2–3 ml of deionized water, and a 55 mm diameter Whatman filter paper (GE Healthcare, WHA1001055) folded into a cone at the bottom of the vial to serve as a moist egg-laying substrate as previously described^52,53^. The females were then dissected to ensure insemination, and the eggs were dried for >24h. The number of eggs per female was counted under the dissection scope using a hand counter. Egg papers were hatched in individual cups, and larvae were counted 4 days later using images captured by a Logitech camera (Logitech, C922x Pro Stream Webcam) and the Cell Counter plugin on FIJI/ImageJ (NIH) as previously described^49^.

#### Insemination and remating assays

##### Spermathecae dissection and scoring

*Spermathecae dissection:* Females were aspirated from their cage and into a paper cup (Solo, KHB16A-J8000) that was covered with a square of 0.8 mm Polyester Mosquito Netting (American Home Habitat, F03A-PONO-MOSQ-M008-WT), secured by two rubber bands (Universal Rubber Bands, Size 12, 0.04” Gauge, Beige, 1 lb Box, 2,500/Pack, UNV00112, Universal) to prevent mosquitoes from escaping. The cup was placed on wet ice for 5 minutes until the females were cold-anesthetized. The top of the cup was tapped intermittently to move the females to the bottom of the cup. 10 ml of 1X PBS (10X PBS buffer, AM9625, Invitrogen; diluted in Millipore water) was pipetted onto a microscope slide (Fisherbrand Superfrost Plus, 12-550-15, Fisher Scientific). Once cold-anesthetized, a single female was placed on the slide next to the 1xPBS drop under a dissection scope (Nikon, Model C-PS), ventral side up. Using Dumont #5 forceps (Roboz Surgical Store, RS-4978), the seventh sternite was gently pressed down, allowing the abdominal tip of the female to extend from the abdomen. With this segment extended, the abdominal tip was easily pulled and removed from the body using forceps and dragged into the 1X PBS drop, and the carcass was discarded from the slide. The spermathecae were then isolated from the remaining tissue, cleaned, and placed on the edge of the drop of 1X PBS so that they could be clearly observed for scoring insemination.

*<u>Scoring insemination</u>*: For insemination scoring, the microscope slide was placed under a compound microscope (SW-350B, SWIFT) with the spermathecae centered in field of view under an air 10x lens with bright field illumination. The female was scored as virgin if the spermathecal sacs were empty and was scored as mated if the spermathecae were full, indicated by sperm presence and darkened color of the sacs. Sperm starts to rotate in the female spermathecae ∼24h after insemination^54^. Using the same methods as for scoring insemination, sperm rotation was examined as an additional verification method for ensuring that previously mated females were indeed mated at the beginning of recording in our behavioral experiments.

*<u>Scoring fluorescent sperm</u>*: Following spermathecae dissection, the slide was placed under a fluorescent microscope (Nikon SMZ18) with the spermathecae centered in the field of view and focus was manually adjusted. The spermathecae were examined under a 10X air lens. Samples were imaged at 475 nm light to score green fluorescent sperm and 549 nm light to score red fluorescent sperm. Fluorescence in both channels was scored as a remated female and no fluorescent sperm was scored as a virgin female. Such females were confirmed to be virgin using the insemination scoring procedure described above.

#### Remating rates after 7 days of co-housing (Figure 1E,F)

60 virgin females (4 days old, wild type) were aspirated into a cage with 30 green-fluorescent sperm males and 30 red-fluorescent sperm males. The animals were left undisturbed for 7 days, after which the females were aspirated from the cage, spermathecae dissected, and sperm color scored.

#### Remating rates at different intervals post-mating (Figure 1G-I)

Per tested interval, 40 virgin females (ages differ per experiment type, see below) were placed in a cage after eclosion. The first male group (red or green-fluorescent sperm) was aspirated into the cage and had access to the females for 30 minutes. At the end of the 30 minutes, the males were aspirated out and discarded. After the specified post mating intervals, the second male group (green or red fluorescent sperm, the other color from the first mating group) were introduced to the cage and had access to the females for 120 minutes. At the end of 120 minutes, the second males were aspirated out of the cage and discarded. The first mating was staggered at intervals prior to a second mating challenge that was held on the same day. The females were dissected the following day and their spermathecae were examined under a fluorescent microscope to score sperm presence and color. Females that did not have the first male sperm color or that only had the second male color, reflecting a failure to mate with the first males, were discarded from the analysis. In each experiment, we also had a group of females that were not introduced to the first males, and were only introduced to the second male group, to verify that successful mating could be obtained under conditions of the second mating (see GitHub^46^ for full results). To avoid any strain-specific biases in mating, we used the green and red sperm males reciprocally as the first male or second male group and we did not detect any differences that arose from the order of introduction (see GitHub^46^ for variation across trials). Two separate experiments were performed to test remating rates at different intervals post mating:

*<u>6 day-old females:</u>* the females were challenged with remating (2^nd^ male group) at six days post eclosion. First matings were staggered prior to the remating challenge and included the examination of the following intervals post initial mating: 2 hours, 6 hours, 24 hours, 48 hours, 72 hours, and no first mating (control group). n=40 females per post-mating interval tested, N=3 trials.

*<u>12 day-old females:</u>* the females were challenged with remating (2^nd^ male group) at twelve days post eclosion. First matings were staggered prior to the remating challenge and included the examination of the following intervals post initial mating: 2 hours, 48 hours, 96 hours, 120 hours, 144 hours, 168 hours, and no first mating (control group). n=40 females per post-mating interval tested, N=3 trials. Per experiment and trial, all intervals were tested in parallel.

#### Behavioral assays and analysis

##### Observing mating attempts

*Comparison of virgin and previously mated females* (**Figure 1A,B**): 15 virgin females and 15 previously mated females (7 days old, wild type) were allowed to acclimate in 30 cm^3^ cages for >6 hours. The cages were then deidentified by another lab member and each cage was examined separately for mating attempts in the following way: 15 males (7 days old, wild type) were aspirated into the cage, followed by a blow of exhaled air to induce flight activity in the cage. The cage was gently tapped every minute to further encourage flight. Mating attempts were scored every 15 seconds by visual observation and indicating the number of mating attempts per timepoint on a spreadsheet. Observation and scoring were carried out for 7 minutes per cage. n=15 pairs per condition tested (virgin or previously mated females), N=3 trials.

*Comparison of intact and manipulated gonostyli males* (**Figure S4C**): 20 5 day-old males were dissected or mock-treated per group. The following day (>24 hours), any dead males were removed from the cage, together with extra males left – to reach a total male amount of 15 males. The cages were deidentified by another lab member and each cage was examined separately for mating attempts in the following way: 15 virgin females (6 days old, wild type) were aspirated into the cage, followed by a blow of exhaled air to induce flight activity in the cage. The cage was gently tapped every minute to further encourage flight. Mating attempts were scored every 15 seconds by visual observation and indicating the number of mating attempts per timepoint on a spreadsheet. Observation and scoring were carried out for 5 minutes per cage. n=15 pairs per condition tested (intact or gonostyli-manipulated males), N=3 trials.

#### 3-dimensional (3D) recordings of mosquitoes in free flight (Figure S2A-I)

*Setup*: A 14 cm^3^ custom clear acrylic cage (see GitHub^46^ for design file) was positioned together with a mirror (10-inch standing cosmetic mirror, DEAYOU, B09XR343HB) angled at 45° to the cage. The camera (sensor: Basler ace acA2040-120um, 35-924; lens: Arducam C-mount Lens for Pi HQ Camera 16mm Focal length, LN045, Arducam) was mounted on a 6-inch stainless steel optical post (Ø1/2 inch Optical Post SS 8-32 Setscrew ¼ inch-20 Tap L = 6 inch 5 Pack, ThorLabs, TR6-P5) for recordings.

*<u>Mosquito handling</u>*: Animals were 4-8 days old at the time of recording. The first animal (female, if recordings were of a male-female pairs) was added to the cage and recording started. After a minute, the second animal (male, if recordings were of a male-female pair) was aspirated into the cage, and the setup was recorded for a total of seven minutes from that point. Exhaled air was blown into the cage through a mouth aspirator (Oral Aspirator with HEPA Filter Model 612, John W. Hock Company) every 30 seconds to encourage flight. Upon completion of the recording, the female was aspirated out of the cage and her spermathecae dissected to score insemination.

*<u>Analysis:</u>* Recorded videos were processed to isolate both the real and reflected views of the behavioral chamber. A custom Python script was used to crop each original video, generating two separate videos for each recording: one for the real view and another for the reflected view (mirror). The trajectories of the mosquitoes in each view were reconstructed separately using two distinct top-down multi-animal classifiers trained in SLEAP (Social LEAP Estimates Animal Poses^55^), one for each view, to detect the position of each mosquito over time. Mosquito identities were verified *a posteriori* by visual inspection, ensuring consistent tracking across frames and between the paired cropped videos derived from the same original recording. After obtaining the tracks for each video, a data integration process was conducted to reconstruct the 3D trajectories. For each video, a scaling factor was determined by the ratio of the vertical position in the real view to the corresponding vertical position in the reflected view for each mosquito. For appropriate 3D reconstruction, we corrected optical distortions by establishing a unified linear relationship between the scaling factor and z-position (i.e. the horizontal position in the reflected view) for each video. This approach accounts for possible aberrations resulting from perspective effects. The resulting 3D trajectories were then interpolated and smoothed to ensure continuity. From the reconstructed 3D trajectories, two parameters were estimated: The Euclidean distance between mosquitoes, and the total distance covered by each mosquito throughout the recording.

### String-tethered female and free flying male interactions (Figure 2, and S2J-L)

*Setup*: A 17.5 cm^3^ polyester and plastic cage (BugDorm Insect Rearing Cage #4S1515) was used for recording, with the clear side facing the camera. A piece of nylon fabric previously worn on arm by a human volunteer was positioned on top of the cage to provide human odor, which enhances flight activity (as described in^10^). Nylon sleeves were prepared by using scissors to remove the fabric from the tip of the stocking foot of knee highs (L’eggs brand Everyday, Amazon), so that the modified stocking was open on both ends^56^. A camera (sensor: Basler ace acA2040-90um, 89-989; lens: Edmund Optics 16mm/ F1.8, 333304) was mounted on a 6-inch stainless steel optical post (Ø1/2-inch Optical Post SS 8-32 Setscrew ¼ inch-20 Tap L = 6 inch 5 Pack, ThorLabs, TR6-P5) for recordings.

*<u>Mosquito handling</u>*: Animals were 4-8 days old at the time of recording. Female mating status was deidentified by a lab member. Females were cold-anesthetized and gently placed on a microscope slide (Fisherbrand Superfrost Plus, 12-550-15, Fisher Scientific) with their dorsal side up. 10 µL of Bondic UV glue (Bondic UV Liquid Plastic Welder, SK8024 BONDIC) was applied to the tip of a 7 cm long clear nylon thread (Clear nylon invisible thread, 017531051647, Blulu) and then attached to the dorsal side of the thorax of the female. UV light (Bondic UV Liquid Plastic Welder, SK8024, BONDIC) was used to cure the glue for 30 seconds. The free end of the string was then attached to the top part of the cage using a reusable glue pad (Uhu Tac Removable and Reusable Glue Pads, SAU99683, UHU) At the start of the recording, the female was prompted to fly by blowing exhaled air into the cage. Once in flight, a male was aspirated into the cage and recording started, and interactions were recorded over an 8-minute period. Throughout the recording, the cage was occasionally tapped gently to stimulate the female’s flight. Females were dissected ∼5 minutes after the completion of recording and their spermathecae examined for the presence of sperm.

*<u>Manual scoring and analysis of behavioral categories:</u>* Recordings were deidentified during the scoring process. Recorded videos were analyzed using BORIS^57^ v. 7.10.7 software. Videos were uploaded to the manual behavioral scoring interface on BORIS and the following behaviors were scored: female flight (state), contact (state), genital contact (state), kicking (state), genital interlocking (state). To accurately define the beginning and end of recordings, “Start” and “End” (events) were scored per video as indicated by the aspiration of the male into and out of the cage, respectively. Binary score of behaviors per second was extracted using the “behaviors binary table” option and plotted as heatmaps on GraphPad Prism (10.4.0). Number of events, average duration, and proportion of total time spent per behavior were extracted using the “synthetic time budget” analysis option and plotted as violins on PRISM.

*<u>Pose estimation analysis:</u>* genital contact and genital interlocking events were initially extracted from the original videos using a custom Python script to process BORIS annotation files. All resulting videos started 2 seconds before the annotation timestamp and lasted until the end of the annotated behavioral event. These were encoded using the open-source FFmpeg library to ensure readability. Next, we trained and implemented a top-down multi-animal pose estimation model in SLEAP^55^ to track five anatomical landmarks per mosquito: the center of the thorax and four sequential points along the abdomen. These points were arbitrarily named, from proximal to distal, abdominal point 1, abdominal point 2, abdominal point 3 and abdominal point 4, with abdominal point 4 representing the abdominal tip. When visible, the lateral characteristic white markings on *Aedes aegypti* abdomens served as reference points for consistent labeling of abdominal segments. After applying the trained model to all processed videos, 181 out of 274 videos met our quality standards for further analysis. This analysis was conducted to determine whether gross differences in the angle formed between the male and female mosquitoes during contact could be informative of their mating status. To characterize the spatial relationship between coupled mosquitoes, we calculated inter-segmental angles between consecutive body parts using the abdominal tips (Abd4) as pivot points. Angle dynamics were analyzed across the entire interaction window to identify potential differences between mating conditions. The following analysis pipeline was developed in Python to process the tracking data: genital contact initiation was defined using temporal and spatial thresholds based on the Euclidean distance between the abdominal tips (Abd4) of paired mosquitoes. We focused our analysis on sustained interactions, defined as continuous genital contact lasting at least 3 seconds (90 frames at 30 frames per second), yielding 174 files out of 181 for final analysis. Videos were categorized into four conditions based on female mating status and interaction type: genital contact in virgin females, genital interlocking, post-genital interlocking genital contact, and genital contact in previously mated females. The results show a consistent pattern across mating conditions with a rapid initial increase from 75° to 90° in the first second followed by a gradual stabilization. No clear statistical differences were observed for the angle formed at the abdominal tips between conditions. Next, we focused on the contribution of male and females to these observed angle dynamics. Because the abdomen of mosquitoes is supple, we decided to analyze the abdominal curvature of both males and females. To quantify abdominal curvature during mating events, we approximated the distal abdominal segment (Abd2-Abd4) using Bezier curves. A Bézier curve is a smooth mathematical curve defined by control points, where the curve passes through the endpoints and is shaped by intermediate control points that influence its curvature. This method allows for flexible fitting of curved trajectories to biological data while maintaining mathematical smoothness and avoiding abrupt directional changes. Sinuosity was calculated as the ratio between the Bezier curve length and the straight-line distance between Abd2 and Abd4. Females maintained lower, more stable sinuosity compared to males, with overlapping confidence intervals across all conditions. Thus, Bezier curve analysis demonstrated minimal variation in abdominal curvature across conditions, indicating that posture alone may not be a reliable predictor of successful mating events. Altogether, these measurements suggest that the basic biomechanical configuration during male-female coupling remains relatively constant regardless of female mating status or interaction outcome.

#### Recordings of pin-tethered female and free flying males interactions in high-resolution

*Setup*: A 14 cm^3^ custom clear acrylic cage was used for recordings (see GitHub^46^ for design file). A camera (sensor: Basler ace acA2040-120um, 35-924; lens: Computar V5024-MPZ C-Mount 50 mm Machine Vision Fixed Focal Lens, BH COV5024MPZ) was mounted on a 6-inch stainless steel optical post (Ø1/2 inch Optical Post SS 8-32 Setscrew ¼ inch-20 Tap L = 6 inch 5 Pack, ThorLabs, TR6-P5), and the focal point was carefully adjusted to focus on the female’s abdominal tip.

*<u>Mosquito handling</u>*: Animals were 4-8 days old at the time of recording. Females were cold-anesthetized and gently placed on a microscope slide (Fisherbrand Superfrost Plus, 12-550-15, Fisher Scientific) with their dorsal side up. 10 ml of Bondic UV glue (Bondic UV Liquid Plastic Welder, SK8024, BONDIC) was applied to the head of a ball head pin (Brass head ball pin, Waydress, B09FGFJKQH) and then attached to the thorax of the female. UV light (Bondic UV Liquid Plastic Welder, SK8024, BONDIC) was used to cure the glue for 30 seconds. The free end of the string was then attached to the top part of the cage using reusable glue pad (Uhu Tac Removable and Reusable Glue Pads, SAU99683, UHU). At the start of the recording, the female was prompted to fly by blowing exhaled air into the cage. Once in flight, 5 males were aspirated into the cage and recording started, and interactions were recorded over a 5-minute period. Throughout the recording, the cage was occasionally tapped gently to stimulate the female’s flight. Females were dissected ∼5 minutes after the completion of recording and their spermathecae examined for the presence of sperm.

*<u>Manual tracking of female dorsal genital tip length and male-female shared surface:</u>* Video recordings of genital interactions of a pin-tethered females and free-flying males were trimmed few seconds before the initiation of genital contact with a male. One genital contact event was tracked per video, selected based on whether genital interlocking took place (virgin females), or the first clearly visible interaction event (previously mated females). Videos were then imported to ImageJ^58^ using the “FFmpeg” import option. The videos were then manually tracked using the “Plugins->Tracking->Manual Tracking” option. Per video, tracking started at the first frame in which the male and female genitalia made contact and tracked for at least 150 frames (on a 37 frames-per-second recording rate), corresponding to 4 seconds. Each track was focusing on a specific point: Track 1= upper end of the female dorsal genital tip. Track 2 = lower end of the female dorsal genital tip. Track 3 = leftmost point of shared contact surface between the male and the female. Track 4 = rightmost point of shared contact surface. Three length measurements were taken of the width of the stem of the metal pin, averaged, and used as a length reference indicating a 0.5 µm real world length. Tracks per sample were then imported to an Excel file and x-y coordinates were used to calculate length of the dorsal genital tip of the female and the shared surface per frame using the following distance formula:

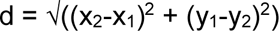

For the average plots, measurements were averaged independently per frame (using the initiation of genital contact as the first aligned frame) and 95% confidence interval was calculated using t-distribution on R (see GitHub^46^).

*<u>Scoring genital interlocking attempts:</u>* Video recordings of pin-tethered females – virgin and previously mated – and free-flying males were examined for instances of genital interlocking attempts by males in the absence of the female genital tip elongation response. Genital interlocking attempts were scored when the male genitalia pushed the female genital tip towards the center of the male genitalia, and additional male genital structures (the paraproct and aedeagus) emerged.

*Female retraction:* To calculate female retraction of dorsal genital tip per behavioral stage, the following measurements were obtained and calculated:

*<u>Tracking pre and post leg contact</u>*: female dorsal genital tip length and male-female shared surface were tracked as described above (“manual tracking”). The first leg contact event was trimmed per video, and each video was then analyzed frame by frame to locate the point in which a leg of the female (any leg) contacted the male. Using this frame as reference, we tracked the female dorsal genital tip and shared surface 75 frames prior to leg contact and 75 frames following leg contact, to track a total of 150 frames in a 37 frames-per-second recording. Per behavioral stage, measurements of 10 frames were extracted and averaged using the following criteria:

*No contact*: the first 10 frames that were measured in the pre-contact tracking. *Leg contact*: the 10 frames surrounding the minimum length of the female dorsal genital tip during leg contact. *Male genital contact*: the first 10 frames that were measured in the male genital contact tracking. *Female acceptance*: the 10 frames surrounding the maximum length of the female dorsal genital tip.

#### Remating Recordings (Figure S3G-K)

*Setup*: A 14 cm^3^ custom clear acrylic cage was used for recordings. A camera (sensor: Basler ace acA2040-120um, 35-924; lens: Computar V5024-MPZ C-Mount 50mm Machine Vision Fixed Focal Lens, BH COV5024MPZ) was mounted on a 6-inch stainless steel optical post (Ø1/2 inch Optical Post SS 8-32 Setscrew ¼ inch-20 Tap L = 6 inch 5 Pack, ThorLabs, TR6-P5) and the focal point was carefully adjusted to focus on the female’s abdominal tip.

*<u>Mosquito handling:</u>* Animals were 4-8 days old at the time of recording. Females were cold-anesthetized and gently placed on a microscope slide (Fisherbrand Superfrost Plus, 12-550-15, Fisher Scientific) with their dorsal side up using forceps (Dumont Tweezers, RS-4978). 10 ml of Bondic UV glue (Bondic UV Liquid Plastic Welder, SK8024, BONDIC) was applied to either string (Clear nylon invisible thread, 017531051647, Blulu) or the head of a ball head pin (Brass head ball pin, Waydress, B09FGFJKQH) and then attached to the thorax of the female. UV light (Bondic UV Liquid Plastic Welder, SK8024, BONDIC) was used to cure the glue for 30 seconds. The free end of the string was then attached to the top part of the cage using reusable glue pad (Uhu Tac Removable and Reusable Glue Pads, SAU99683, UHU). At the start of the recording, the female was prompted to fly by blowing exhaled air into the cage. Once in flight, 5 green-sperm males were aspirated into the cage and recording started. The recording was monitored in real-time to detect female acceptance and genital interlocking. Once genital interlocking and mating occurred and the male detached from the female, the green-sperm males were quickly aspirated out of the cage and 5 red-sperm males were immediately aspirated into the cage and interactions were recorded over an additional 5-minute period. If mating did not take place, the green-sperm males were removed from the cage after 5 minutes of recording and replaced with the red-sperm males for an additional 5 minutes of recording. Throughout the recording, the female was stimulated to fly by occasional puffs of exhaled air. Females were dissected ∼10 minutes after the completion of recording and their spermathecae examined under a fluorescent microscope to score presence and color of sperm.

#### Pin-tethered male and female genital interaction recordings of same-species and cross-species genital interactions (Figure 7)

*<u>Setup</u>*: A dissection microscope (Nikon SMZ645), with the eyepieces removed and replaced with the iDuOptics iPhone mount (LabCam for iPhone, iDu Optics, B0799QTHVR) and an iPhone X attached. The built-in recording software was used, recording at 30 frames per second.

*<u>Mosquito handling:</u>* Females and males were collected separately in net-covered paper cups (Solo, KHB16A-J8000) and cold-anesthetized. The female was gently placed on a microscope slide (Fisherbrand Superfrost Plus, 12-550-15, Fisher Scientific) with her dorsal side up using forceps (Dumont Tweezers, RS-4978). 10 ml of Bondic UV glue (Bondic UV Liquid Plastic Welder, SK8024, BONDIC) was applied to the head of a ball head pin (Brass head ball pin, Waydress, B09FGFJKQH) and then attached to the thorax of the female. UV light (Bondic UV Liquid Plastic Welder, SK8024, BONDIC) was used to cure the glue for 30 seconds. The female was then tethered perpendicular and elevated 5 cm above the dissection scope surface using a reusable glue pad (Uhu Tac Removable and Reusable Glue Pads, SAU99683, UHU). The orientation of the female was confirmed through the iPhone X so that her lateral left side was being observed, with her ventral side facing up in the recording. The male was similarly treated with the UV glue and a ball head pin but was not tethered so that his position could be controlled and adjusted by the experimenter. The tethered female was brought into focus and her terminalia was centered in the field of view of the iPhone X. The video recording was started and the experimenter, by holding the edge of the male’s metal pin by hand, gently positioned the male so that its abdominal tip touches the female tip and the pair is in genital contact. The process did not involve forceful pushing and instead the abdominal tips of the pair were gently “brushed” against each other. Throughout the recording, we ensured that the female’s terminalia was consistently in focus by manually adjusting the focus. Recordings lasted until genital interlocking was complete or until the 8-minute mark was reached. The female was allowed to recover for 15 minutes while still tethered to the metal pin before her spermathecae were dissected and scored for <u>the presence of sperm</u>.

*<u>Scoring behaviors in pin-tethered male and female genital interaction recordings:</u>* Videos were deidentified and each video was individually scored for the three characterized behaviors: 1) male genital contact, 2) female genital tip elongation, and 3) genital interlocking. Since the position of the male was controlled in this experimental setup, male genital contact was scored as a postural change of the male tip aimed towards a tighter connection with the female tip, and the rapid vibration of the gonostyli when present. Female genital tip elongation was scored as extension of the female genital tip from its resting or retracted position. Genital interlocking was scored when the male and female formed a tight connection of their genitalia, and was further verified by gently pulling the male away from the female to check that the animals do not separate and remain interlocked.

#### Gonostylus manipulation assays

##### Surgical manipulation of the gonostylus (Figure 5 and Figure S4A-B)

*<u>General procedure</u>*: 3-day old males were cold-anesthetized on ice and were gently placed on a microscope slide (Fisherbrand Superfrost Plus, 12-550-15, Fisher Scientific) ventral-side up under a dissection scope (Nikon SMZ645). Focusing in on the genital tip of the male, forceps (Dumont Tweezers, RS-4978) were used to gently extend the gonostyli from their resting position and make the following manipulations:

*Mock treatment*: Males were cold-anesthetized for 15 minutes and placed in a paper cup (Solo, KHB16A-J8000) for recovery.

*Removed gonostyli:* Forceps were used to pinch at the base of the gonostylus and remove the entire structure, with minimal invasiveness of the hairs.

*Truncated gonostyli:* Forceps were used to pinch at the half-length mark of the gonostyli and remove the distal end of the structure.

*<u>Claws removed:</u>* Forceps were used to pinch at the tip of the gonostyli to remove the claws. *<u>Dislocation (immobilize the structures):</u>* Forceps were used to gently press alongside the base of the gonostyli resulting in lack of muscle control to the gonostyli and their immobilization

Following the surgical manipulations, males were placed in a paper cup (Solo, KHB16A-J8000) and allowed to recover for 24 hours in a 30cm^3^ BugDorm-1 Insect Rearing Cage (MegaView Science). 24 hours after the manipulation the males were checked for their ability to fly by blowing exhaled air into the cage, and the paper cup was checked to see if any males did not survive the procedure.

#### Insemination success of males with manipulated gonostyli

Per manipulation condition, 30 virgin females were placed into the cage of 30 males – all with the same gonostyli manipulation – which were added 24 hours prior for their post-manipulation recovery. The animals were left undisturbed for 48 hours to allow ample time for mating. At the end of the 48-hour period, females were aspirated out of the cage and their spermathecae were scored for sperm presence. N=4 trials.

*<u>Competition success:</u>* using the two transgenic strains, green-fluorescent and red fluorescent sperm males, we conducted mating competition assays between males with manipulated and intact (mock treated) gonostyli. 20 virgin females (4 days old, wild type) were placed in a 30 cm^3^ cage, to which 15 manipulated and 15 intact males were added. Per replicate and tested manipulation, a reciprocal use of the two sperm color strains was carried out to prevent strain biases in competitive mating. For each replicate one tested group had the red sperm males manipulated and the greens intact, and another group had the green sperm males manipulated (same manipulation) and the red intact. The males and females were co-housed for 48 hours, after which the females were aspirated out and their spermathecae were dissected and examined under a fluorescent microscope to score sperm presence and color. Females which had two sperm colors present (remating) were omitted from the analysis. Full results are available on GitHub^46^.

#### Scanning Electron Microscopy

##### Conventional scanning electron microscopy

7-day old male and virgin female mosquitoes were fixed in 2% glutaraldehyde, 4% paraformaldehyde, in 0.1M sodium cacodylate buffer pH 7.2 (0.4M buffer - Electron Microscopy Sciences, 11655-1G - diluted with Millipore water), for 3 hours at room temperature. After the initial fixation, the terminus of the female abdomen was severed with a razor blade between the 4th-5th abdominal segments, and the anterior portions were discarded. Fixation of the abdominal tip continued for 2 hours at room temperature and then overnight at 4°C. After rinsing three times with cacodylate buffer, samples were dehydrated in a graded series of ethanol (30%, 50%, 70%, 90%) for 10 minutes each and three times in 100% ethanol with molecular sieves for 15 minutes each and then critical point dried in a Tousimis Autosamdri 931. After mounting, samples were sputter coated with 10 nm of iridium with a Leica ACE600 coater. Samples were imaged on a JEOL JSM-IT500HR scanning electron microscope at 5.0 kV. Adobe Illustrator was used to generate falsely colored SEM images.

##### Cryo-scanning electron microscopy

30 7-day old virgin females were placed in a 30 cm^3^ cage. 30 males were then aspirated to the cage and the mosquitoes were carefully observed by eye to detect genital interlocking events. Upon detection of a pair in genital interlocking, the experimenter quickly aspirated the couple using a mouth aspirator (Oral Aspirator with HEPA Filter Model 612, John W. Hock Company) directly into liquid nitrogen. The frozen couple was mounted on a cryo-stub (JEOL, Cat. JU2010535) precooled at liquid nitrogen temperature. Samples were imaged within 30 minutes on a JEOL JSM-IT500HR scanning electron microscope under low vacuum mode (30 Pa of pressure) at 20 KV.

### QUANTIFICATION AND STATISTICAL ANALYSIS

#### Recording software

BASLER video recording software was used on a Dell Latitude 14 Rugged laptop (Model type: P46G001) to capture the recordings for the 3D, string-tethered, pin-tethered, and remating behavioral assays.

#### Code and analysis plots

Plots created in this study were generated using Python (3.11), GraphPad Prism (10.4.0), and R (3.6.1). Code development was assisted by Claude (Anthropic, 2024-2025), with all code being manually reviewed and validated by the authors. Details and code are available at GitHub (https://github.com/VosshallLab/HouriZeevi_Vosshall-2025).

### ADDITIONAL RESOURCES

Data and analysis code used in this publication, as well as Extended data Video 1 (comprising 10 individual videos), are available at Zenodo (https://zenodo.org/records/15133603) and GitHub (https://github.com/VosshallLab/HouriZeevi_Vosshall-2025).

## Supplemental Figures

**Figure S1:**
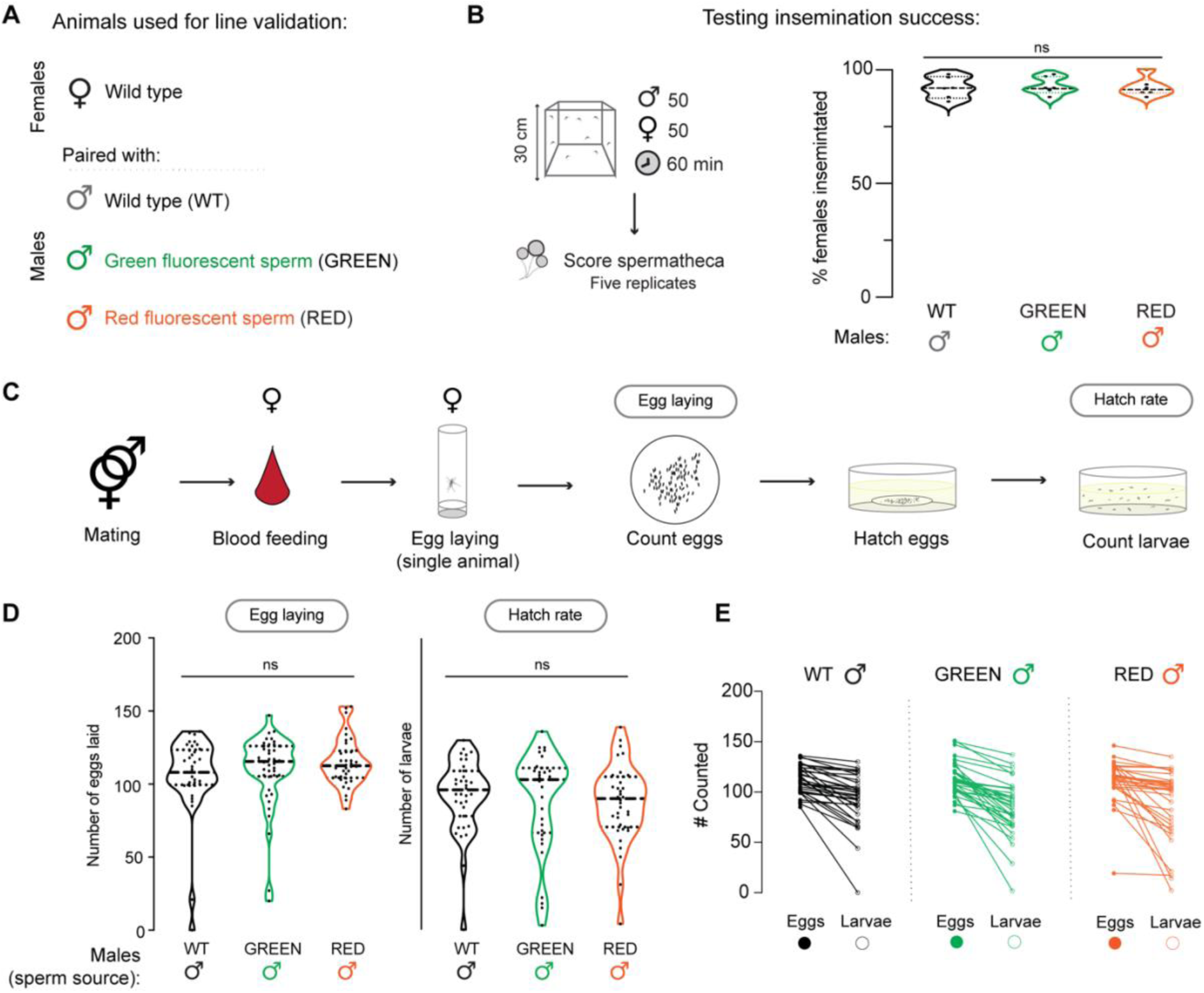
Testing mating success and fertility of transgenic fluorescent sperm males. Related to Figure 1. **A**. Animals used in **B-E**. **B**. Schematic of experimental setup (left) and results (right) for testing insemination rates of transgenic males. N=5 trials with 50 females and 50 males each. Median (thick dotted line) and 25/75 quartiles (thin dotted line) are indicated. Data were analyzed using the Mann-Whitney test. ns=not significant. **C**. Schematic of experimental setup for testing egg laying and hatch rate. **D,E**. Results for egg laying and hatch rate, (**D**) plotted separately (median – thick dotted line – and 25/27 quartiles – thin dotted line – indicated) or (**E**) paired. n=36-40 eggs/larvae obtained from single females were tested per condition. Data were analyzed using the Mann-Whitney test. ns=not significant.

**Figure S2:**
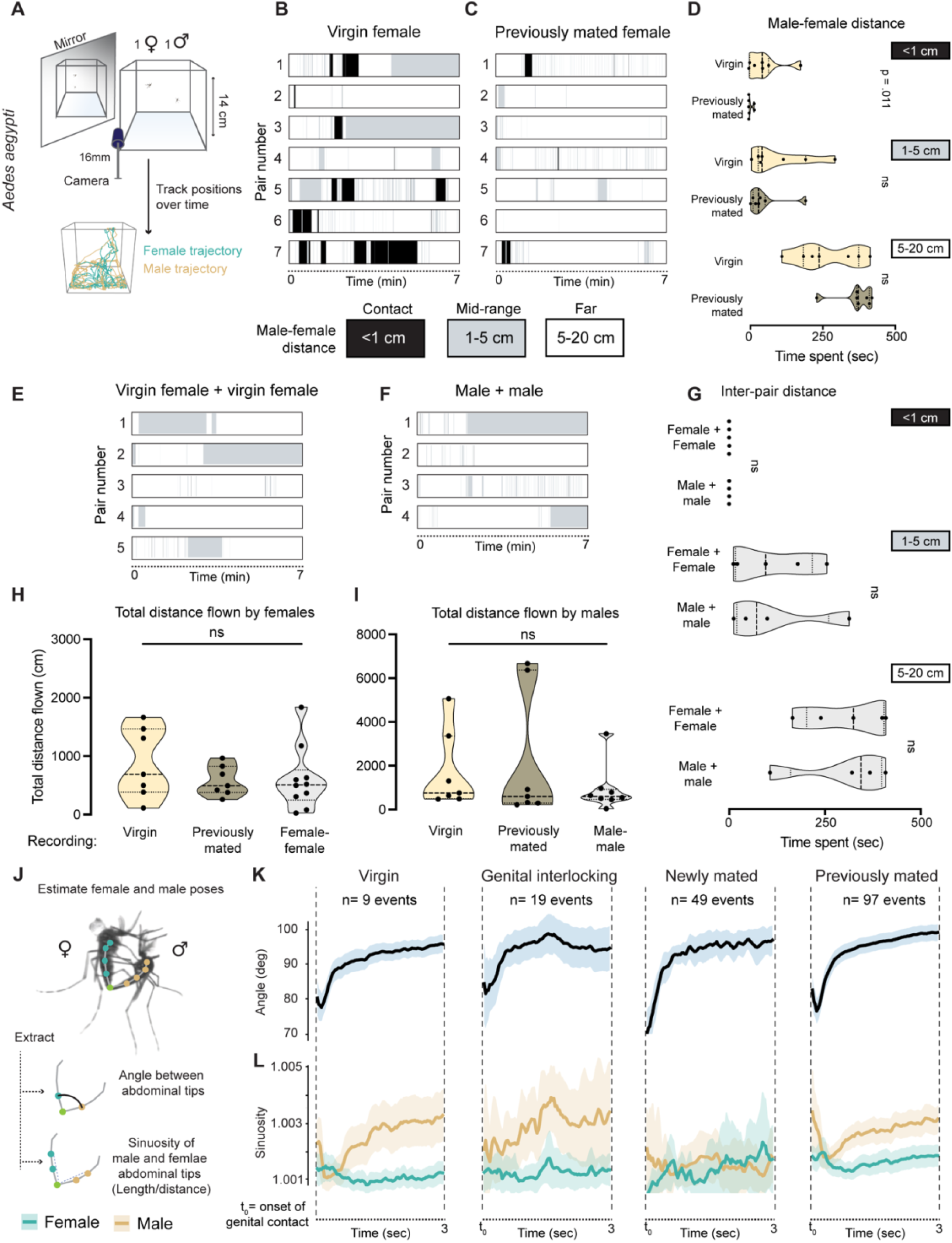
Interactions of free flying mosquito pairs (distance) and string-tethered females and free-flying males (pose estimate). Related to Figure 2. **A**. Schematic of experimental setup for 3-dimensional recording of free flying male and female mosquitoes (1 each) (top) and example of the male and female trajectories over the course of the 7-minute recording (bottom). **B,C.** Raster of distance between pairs of mosquitoes in (**B**) virgin female and (**C**) previously mated female recordings. Virgin females #2 and #4 were not inseminated after 7 minutes. N=7 recordings per condition. **D.** Summary of time spent in the different distance categories: contact (<1 cm), mid-range (1-5 cm), and far (5-20 cm). Median (thick dotted line) and 25/75 quartiles (thin dotted line) are indicated. Data were analyzed using the Mann-Whitney test. ns=not significant. **E, F**. Distance analysis of pairs of (**E**) virgin females or (**F**) males. N=5 (**E**) and N=4 recordings (**F**). **G.** Summary of time spent in the different distance categories: contact (<1 cm), mid-range (1-5 cm), and far (5-20 cm). Median (thick dotted line) and 25/75 quartiles (thin dotted line) are indicated. Data were analyzed using the Mann-Whitney test. ns=not significant. **H,I** Summary of total distance flown by (**H**) females and (**I**) males across all recordings. Median (thick dotted black line) and 25/75 quartiles (thin dotted black line) are indicated on violin plots. N=7 animals plotted for virgin and previously mated recordings. (**H**) N=10 and (**I**) N=8. Data were analyzed using the Mann-Whitney test. ns=not significant. **J.** Analysis setup of pose estimation using SLEAP ^S1^ during genital contact events in string-tethered female recordings. **K,L.** Dynamics of the first 3 seconds of genital contact events, showing the (**K**) shared angle between the male and the female, and the (**L**) sinuosity of the male (tan) and female (turquoise) plotted separately for the indicated event types.

**Figure S3:**
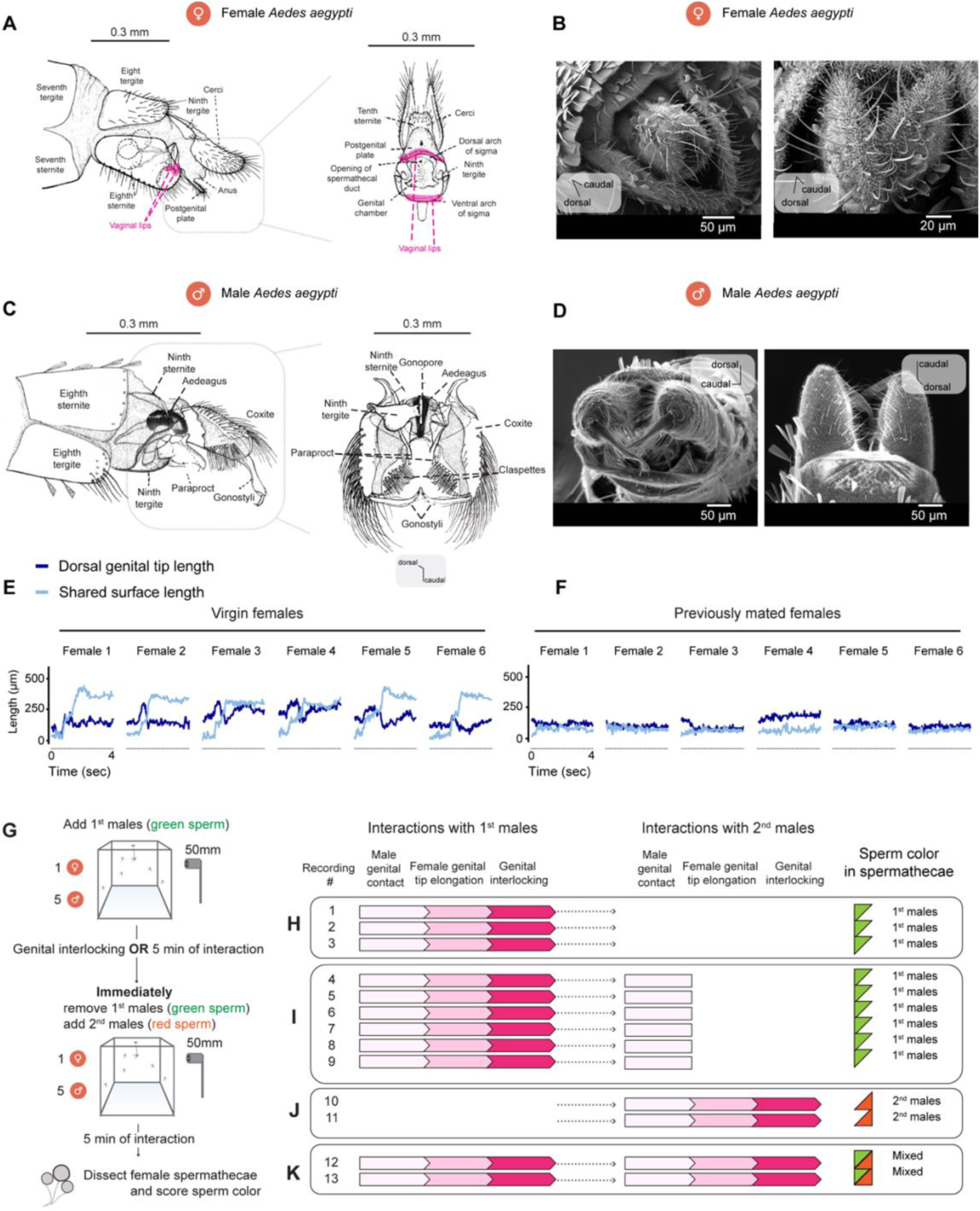
Female and male genital anatomy and female genital tip elongation as the gating mechanism for mating and remating. Related to Figure 3. **A.** Reproductions of anatomical drawings female *Aedes aegypti* genital structures (after Jobling, 1987) ^S2^. Female tip is presented in a lateral view (left) and caudal view with the vaginal lips spread open (right). Vaginal lips are colored pink. **B**. Scanning electron microscopy of female tip. **C**. Reproductions of anatomical drawings male *Aedes aegypti* genital structures (after Jobling, 1987)^S2^. Male tip is presented in a lateral view (left) and ventral view (right). **D**. Scanning electron microscopy of male tip. **E,F**. Individual tracks of female dorsal genital tip length and the male-female shared surface length during genital interactions of (**E**) virgin and (**F**) previously mated females. **G**. Schematic of experimental setup for recording genital behavior and scoring sperm color during sequential mating. **H-K**. Genital behaviors and post-recording scoring of sperm color in spermathecae. Each row corresponds to a recording of a single female. N=13 recordings with 1 female and two groups of 5 males each.

**Figure S4:**
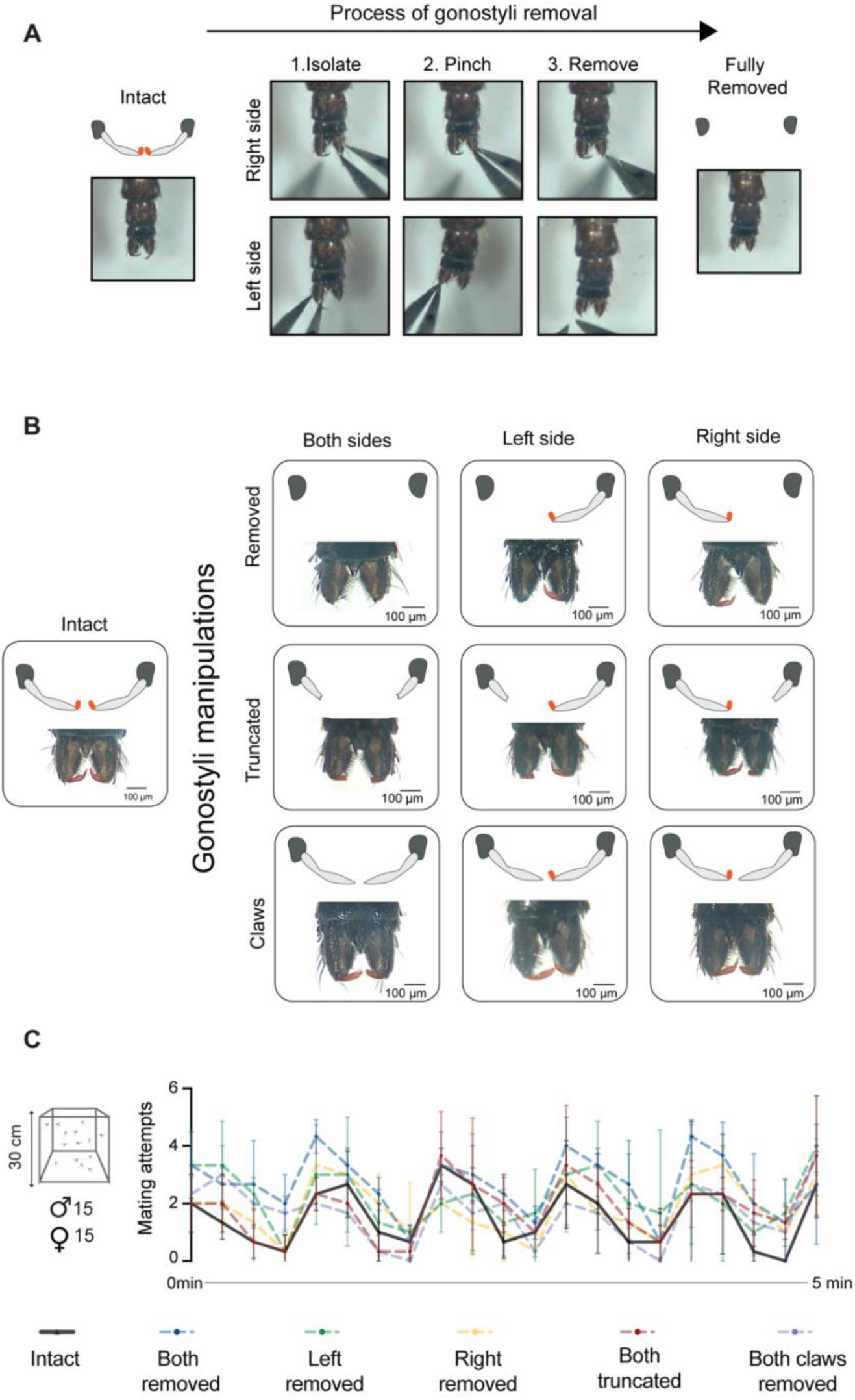
Gonostyli manipulation and mating attempts of intact and gonostyli-manipulated males. Related to Figure 5. **A**. The step-by-step procedure for removing the gonostyli. **B**. Representative snapshots of intact gonostyli and the different gonostyli manipulations. Gonostyli are pseudo-colored orange. **C**. Mating attempts observed for 5 minutes in cages of virgin females and intact or gonostyli-manipulated males. N=3 trials per male condition, with 15 males and 15 virgin females each. Cages were gently tapped every minute to induce flight activity, which induces a corresponding increase in mating attempts. Black line (intact) and dotted lines (males with the indicated gonostylus manipulation, color coded in legend at bottom) connect the mean mating attempts per indicated timepoint. Error bars represent standard deviation.

**Figure S5:**
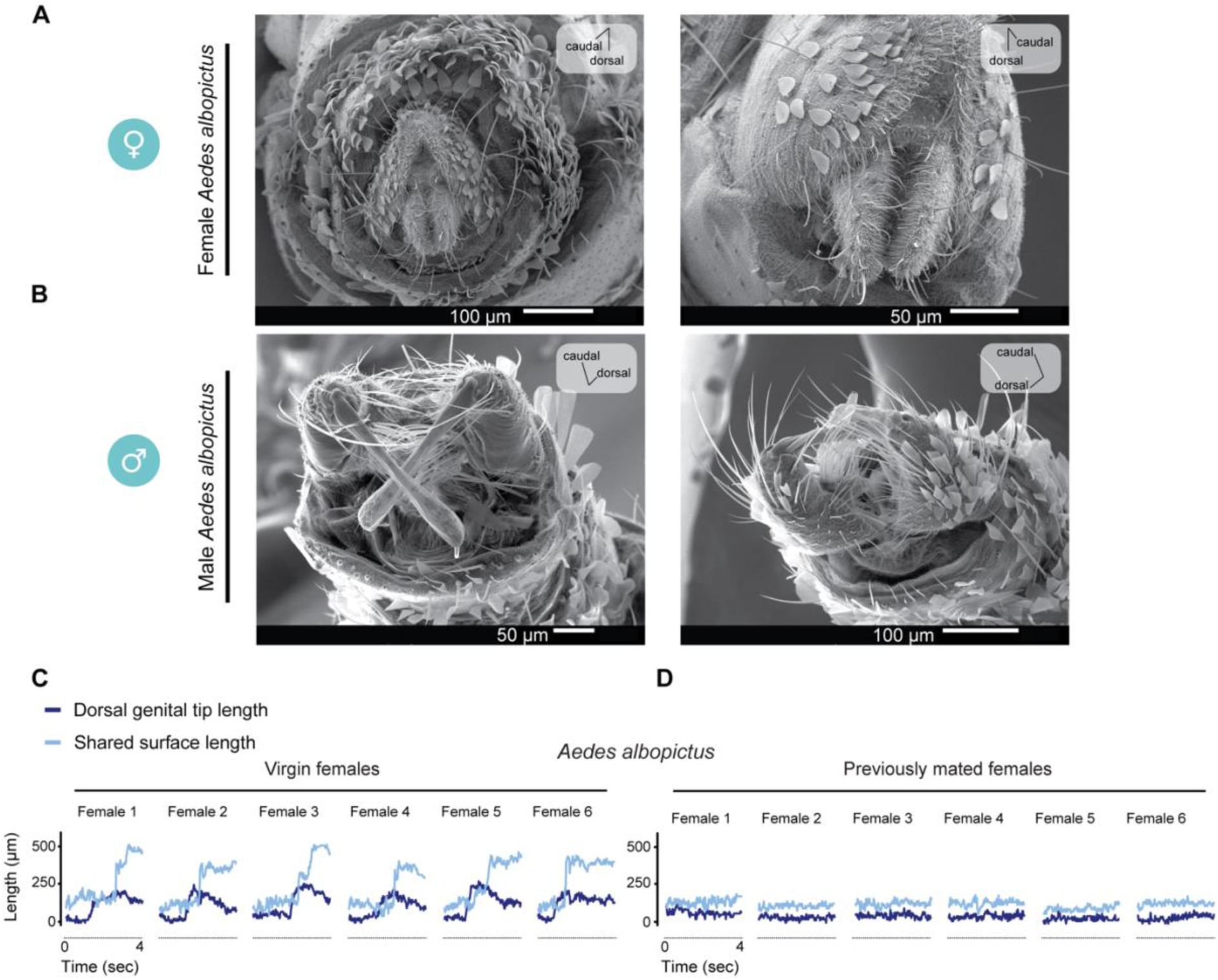
*Aedes albopictus* genital anatomy and genital interaction dynamics. Related to Figure 6. **A,B.** Scanning electron microscopy images of (**A**) female and (**B**) male genitalia of *Aedes albopictus*. **C,D**. Individual tracks of female dorsal genital tip and male-female shared surface dynamics during genital interactions in *Aedes albopictus* from recordings of (**C**) virgin females or (**D**) previously mated females.

